# Delta-Omicron recombinant escapes therapeutic antibody neutralization

**DOI:** 10.1101/2022.04.06.487325

**Authors:** Ralf Duerr, Hao Zhou, Takuya Tada, Dacia Dimartino, Christian Marier, Paul Zappile, Guiqing Wang, Jonathan Plitnick, Sara B. Griesemer, Roxanne Girardin, Jessica Machowski, Sean Bialosuknia, Erica Lasek-Nesselquist, Samuel L. Hong, Guy Baele, Meike Dittmann, Mila B. Ortigoza, Prithiv J. Prasad, Kathleen McDonough, Nathaniel R. Landau, Kirsten St. George, Adriana Heguy

**Author notes:** Co-corresponding authors **Corresponding authors** Ralf Duerr, MD, PhD, Department of Microbiology, NYU Grossman School of Medicine, Alexandria Center for Life Science (ACLS), West Tower, 430 East, 29th Street, Room 323 New York, NY 10016, Adriana Heguy, PhD, Department of Pathology, NYU Grossman School of Medicine, Genome Technology Center, NYU Langone Health, 550 First Avenue, MSB 294A, New York, NY 10018. Shared contribution (Ralf Duerr, Hao Zhou, Takuya Tada).

## Abstract

**Background:** The emergence of recombinant viruses is a threat to public health. Recombination of viral variants may combine variant-specific features that together catalyze viral escape from treatment or immunity. The selective advantages of recombinant SARS-CoV-2 isolates over their parental lineages remain unknown.

**Methods:** Multi-method amplicon and metagenomic sequencing of a clinical swab and the *in vitro* grown virus allowed for high-confidence detection of a novel recombinant variant. Mutational, phylogeographic, and structural analyses determined features of the recombinant genome and spike protein. Neutralization assays using infectious as well as pseudotyped viruses and point mutants thereof defined the recombinant’s sensitivity to a panel of monoclonal antibodies and sera from vaccinated and/or convalescent individuals.

**Results:** A novel Delta-Omicron SARS-CoV-2 recombinant was identified in an unvaccinated, immunosuppressed kidney transplant recipient treated with monoclonal antibody Sotrovimab. The recombination breakpoint is located in the spike N-terminal domain, adjacent to the Sotrovimab quaternary binding site, and results in a 5’-Delta AY.45 and a 3’-Omicron BA.1 mosaic spike protein. Delta and BA.1 are sensitive to Sotrovimab neutralization, whereas the Delta-Omicron recombinant is highly resistant to Sotrovimab, both with and without the RBD resistance mutation E340D.

**Conclusions:** Recombination between circulating SARS-CoV-2 variants can functionally contribute to immune escape. It is critical to validate phenotypes of mosaic viruses and monitor immunosuppressed COVID-19 patients treated with monoclonal antibodies for the selection of recombinant and immune escape variants. (Funded by NYU, the National Institutes of Health, and others)

## Introduction

In individuals infected with more than one viral variant, template switching during viral RNA synthesis can lead to RNA recombination, resulting in mosaic viruses^1^. As known from influenza and HIV, recombination can accelerate viral adaptive evolution^2,3^. Recombination may result in viruses that have increased transmissibility or evade the host immune response or antiviral treatment. However, causal links between recombination events and functional selective advantages have not yet been described for SARS-CoV-2. Recombination events have been recorded in seasonal coronaviruses and MERS-CoV, and are discussed as cause for the initial zoonotic spillover of SARS-CoV-2^4^. Within the first year of the SARS-CoV-2 pandemic, there was little evidence of recombination, similar to the short-lived SARS-CoV-1 epidemic^5^. Since then, high case numbers and the co-circulation of variants have increased the likelihood of coinfections^6,7^ and the opportunity for inter-variant recombination^6,8-13^.

Delta was the dominant variant in the second half of 2021 and remained in circulation through early 2022. Omicron has caused a novel global wave beginning in November 2021, which overlapped with Delta for several weeks^14^. In New York City (NYC), Omicron arrived when the only circulating variant was Delta and its sublineages^15^. Some Delta-Omicron recombinants were identified in Europe and the U.S.^6,8,12^. In these recombinants, genome break points frequently occurred in the N-terminal domain (NTD) of spike, resulting in hybrid genomes where non-structural ORF1a/b genes are from Delta and C-terminal spike regions, including the receptor-binding domain (RBD) from Omicron. Data on the functional consequences of recombination are scarce. Two studies using pseudotyped virus neutralization or recombinant spike infectious reporter assays suggested that a Delta-Omicron recombinant had resistance comparable to BA.1 with regards to convalescent or vaccine-induced antibodies^16,17^. The sensitivity to monoclonal antibodies (mAbs) was not tested, though mAbs are a key component to treat immunocompromised patients, in whom new variants often emerge. Overall, there is a lack of data on which underlying conditions and treatment pressures favor the selection of recombinant SARS-CoV-2 over their parental lineages.

Here, we report a novel Delta (AY.45)-Omicron (BA.1) recombinant, detected in an immunosuppressed, unvaccinated COVID-19 patient who was treated with Sotrovimab, a lead nAb against the BA.1 Omicron lineage. We show that, whereas Delta and BA.1 are sensitive, the recombinant virus is resistant to neutralization by Sotrovimab, irrespective of the presence of a rare E340D resistance mutation in the spike RBD. To our knowledge, this is the first report of a recombinant SARS-CoV-2 spike conferring escape from a therapeutic mAb regimen.

## Methods

Detailed methods can be found in the Supplementary Appendix.

### SARS-CoV-2 sequencing and bioinformatic analysis

SARS-CoV-2 full-genome sequencing was performed in two independent laboratories using four different approaches to confirm the identity of the recombinant and to exclude artifacts (e.g. by mixed reads) as reason for the detection of recombinant sequences. The methods included xGen Amplicon^18^ and metagenomics sequencing (NYU), as well as AmpliSeq Insight and ARTIC Amplicon sequencing (NY State DOH).

### Virus isolation and culture

VeroE6/TMPRSS2 cells^19^ were used for virus isolation, obtained from the Japanese Collection of Research Bioresources (JCRB Cell Bank) cell number JCRB1819, through Sekisui Xenotech, LLC (Kansas City, KS), agreement # A2000230. VeroE6/TMPRSS2 cells were seeded three days prior to infection, to reach 85-90% confluency. Infected monolayers were checked daily for cytopathic effect (CPE), and 110 µL of supernatant was removed at 24, 48, 72, and 96 hours post-infection (hpi) for further analysis. At 96 hpi, cells and supernatant were harvested together.

### Mutation and phylogenetic analysis

Highlighter analyses were performed on MAFFT-aligned SARS-CoV-2 full-genome sequences using the Highlighter tool provided by the Los Alamos HIV sequence database^20^. Phylogenetic analyses were performed using the Nextstrain CLI package using a New York, New Jersey, and Connecticut-focused subsampling of global SARS-CoV-2 sequences^14^.

### Phylogeographic analysis

Bayesian phylogeographic analysis were performed on the Delta (AY.45) piece of the recombinant using BEAST v1.10.5^21^.

### Structural analysis

Molecular graphics and analyses were performed with UCSF ChimeraX 1.3^22^. Homology models of the recombinant spike were generated with the protein structure homology-modelling server SWISS-MODEL^23^.

### Infectious virus neutralization assay

Plaque reduction neutralization tests (PRNTs) were performed using low passage, sequence-confirmed SARS-CoV-2 virus isolates containing approximately 100-180 plaque forming units and 2-fold serially diluted monoclonal antibody preparations or test sera on confluent Vero E6 with TMPRSS2 cells (JCRB1819, Sekisui XenoTech). IC50 values were calculated by determining the percent neutralization (relative to virus only controls) for technical duplicates of each well and using non-linear regression (GraphPad Prism).

### Pseudotyped virus neutralization assay

The SARS-CoV-2 BA.1-spike was previously generated^24^. The N-terminal domain (Delta-like) of the SARS-CoV-2 Delta-Omicron recombinant spike was chemically synthesized as a short fragment (Genscript) and fused by overlapping PCR with the RBD and C-terminal parts of the BA.1 spike. The full Delta-Omicron recombinant spike was cloned into pcDNA6 (Invitrogen). Point mutations were introduced by overlap extension PCR. All spike expression vectors encode a termination codon that deletes the carboxy-terminal 19 amino acids (Δ19)^25^. HIV-1 Gag/Pol expression vector pMDL and HIV-1 Rev expression vector pRSV.Rev have been previously described^25^. SARS-CoV-2 variant spike pseudotyped lentivirus stocks were produced by cotransfection of 293T cells with pMDL, pLenti.GFP-NLuc, pcCoV2.S-Δ19 (or variants thereof) and pRSV.Rev^25^. Neutralizing titers were determined using serially diluted sera or mAbs on ACE2.293T cells 1 day after pseudotyped virus transduction (MOI 0.2). D614G virus was used as reference. Experiments were performed in technical duplicates and statistical significance determined by two-tailed, unpaired t-tests or one-way ANOVA (*P≤0.05, **P≤0.01, ***P≤0.001, ****P≤0.0001, GraphPad Prism 8). Confidence intervals are shown as the mean ± standard deviation.

### Human sera

Sera from individuals vaccinated at NYULH with BNT162b2 were collected through the NYU Vaccine Center with written consent under I.R.B. approval and were deidentified.

### Study approval

This study was approved by the NYULH Institutional Review Board, protocol numbers i21-00493, i21-00561, i20-00595, and i18-02037 (NYU Langone Vaccine Center).

## Results

### An immunosuppressed transplant recipient with two COVID-19 episodes, treated with Sotrovimab

A male in his late twenties had end-stage renal disease due to hypertensive nephrosclerosis and received a deceased donor kidney transplant in June 2021. He was unvaccinated against COVID-19. His immunosuppressive regimen consisted of thymoglobulin induction at transplant and tacrolimus, prednisone, and mycophenolate mofetil as maintenance (**Table 1**).

**Table 1.** Demographic and clinical features of a patient infected with a Delta-Omicron recombinant SARS-CoV-2 virus.

| Demographics |  |
| --- | --- |
| Age range | 25-29 |
| Sex | Male |
| Clinical features |  |
| COVID-19 vaccination status | Unvaccinated |
| Immune status | Immunosuppressed |
| Comorbidities | Hypertension, |
|  | End-stage renal disease (ESRD) due to hypertensive nephrosclerosis |
|  | Deceased donor kidney transplant recipient in 2021 |
| Immunosuppression at baseline | Induction (at transplant): Thymoglobulin |
| Maintenance immunosuppression | Tacrolimus 2mg q AM / 3 mg q PM, prednisone 5 mg daily, mycophenolate mofetil 1000 mg q12h |
| COVID-19 clinical course |  |
| 1) December 2021 | COVID-19 positive by rapid antigen test (no PCR done) |
|  | Cough and fever |
|  | Not hospitalized |
|  | Received Sotrovimab 500 mg infusion once |
|  | Symptoms resolved the next day |
| 2) February 2022 | Fever (up to 102°F), chills, body aches, and fatigue for 4 days |
|  | Hospitalized |
|  | No cough, shortness of breath, hypoxia, or chest pain. Not on supplemental oxygen |
|  | Chest X ray: right basilar opacities |
|  | Chest CT: left lower lobe pneumonia |
| Tests for potential pathogens | Respiratory viral pathogen panel multiplex<br>PCR positive for SARS-CoV-2 (Ct 26.4) |
|  | Negative for blood cultures, *Legionella* and *Streptococcus pneumoniae* urinary antigens, serum cryptococcal antigen, beta-D- glucan, and Aspergillus galactomannan. |
|  | Sputum culture was not performed due to lack of a good quality specimen. |
| Treatment | No antivirals given |
|  | Antibiotics for 7 days (2 days Piperacillin-tazobactam + vancomycin, then 2 days piperacillin-tazobactam + azithromycin, then 3 days Levofloxacin) |
|  | Immunosuppression reduced by halving mycophenolate dose, and maintaining the doses of tacrolimus and prednisone |
| Outcome | Afebrile after 2 days of antibiotic treatment |
|  | Discharged after 4 days |

#### 1^st^ COVID-19 episode

In late December 2021, he developed cough and fever and tested positive for COVID-19 by rapid antigen test on the same day. No PCR test was done. He received an infusion of Sotrovimab 500 mg two days after onset of symptoms which resolved the next day, and he did not require hospitalization at this time.

#### 2^nd^ COVID-19 episode

In the middle of February 2022, he presented with a fever up to 102^°^F, chills, body aches, and severe fatigue for one day, prompting hospitalization. He had no cough, shortness of breath, or chest pain. A chest X-ray showed left-sided basilar opacities. A CT scan of the chest showed patchy airspace opacities with consolidation in the left lower lobe consistent with pneumonia. A respiratory viral pathogen panel multiplex PCR on a nasopharyngeal swab specimen was positive for SARS-CoV-2 (Ct 26.4). He received empiric antibiotics for a presumed bacterial superinfection for seven days, and his immunosuppressive regimen was modified by halving the dose of mycophenolate. Tacrolimus and prednisone were continued at previous doses. The patient became afebrile two days into hospitalization, and was discharged two days later. He did not receive antiviral medications during or after hospitalization. No further RT-PCR test was done and no hypoxia was observed (**Table 1**).

### Identification of a Delta-Omicron recombinant virus and phylogeographic analysis

Genomic surveillance in the greater New York City area revealed an unusual SARS-CoV-2 sequence in late February 2022, based on its outlier Omicron BA.1 placement in a global phylogenetic tree and its high number of mutations (**Figure 1A,B**). Pangolin did not assign a lineage^26^, and further inspection of each individual mutation and comparison with the mutations from different lineages revealed that the entire ORF1ab genomic region and 5’region of the spike gene up to position 22,035 contained Delta-specific mutations most similar to Delta sublineage AY.45 and lacked Omicron-specific mutations. The remainder of the genome, specifically after position 22,193 contained Omicron BA.1-specific mutations and had no Delta-specific mutations, indicative of a Delta-Omicron recombinant with a single breakpoint in spike NTD (**Figure 1A**). Phylogenetic analyses of the separate subregions confirmed the relatedness of the 5’ and 3’ segments with Delta (AY.45) and Omicron (BA.1) clades, respectively (**Figure 1B**). Nine multi-method sequencing runs on different specimens, including xGen amplicon sequencing, metagenomics, Ion AmpliSeq Insight, and *in vitro* virus growth with ARTIC sequencing confirmed the identity of a Delta-Omicron recombinant (**Table S1**; **Supplemental Results**). The sequence was deposited in GISAID (EPI_ISL_10792641), flagged as a recombinant.

**Figure 1.**
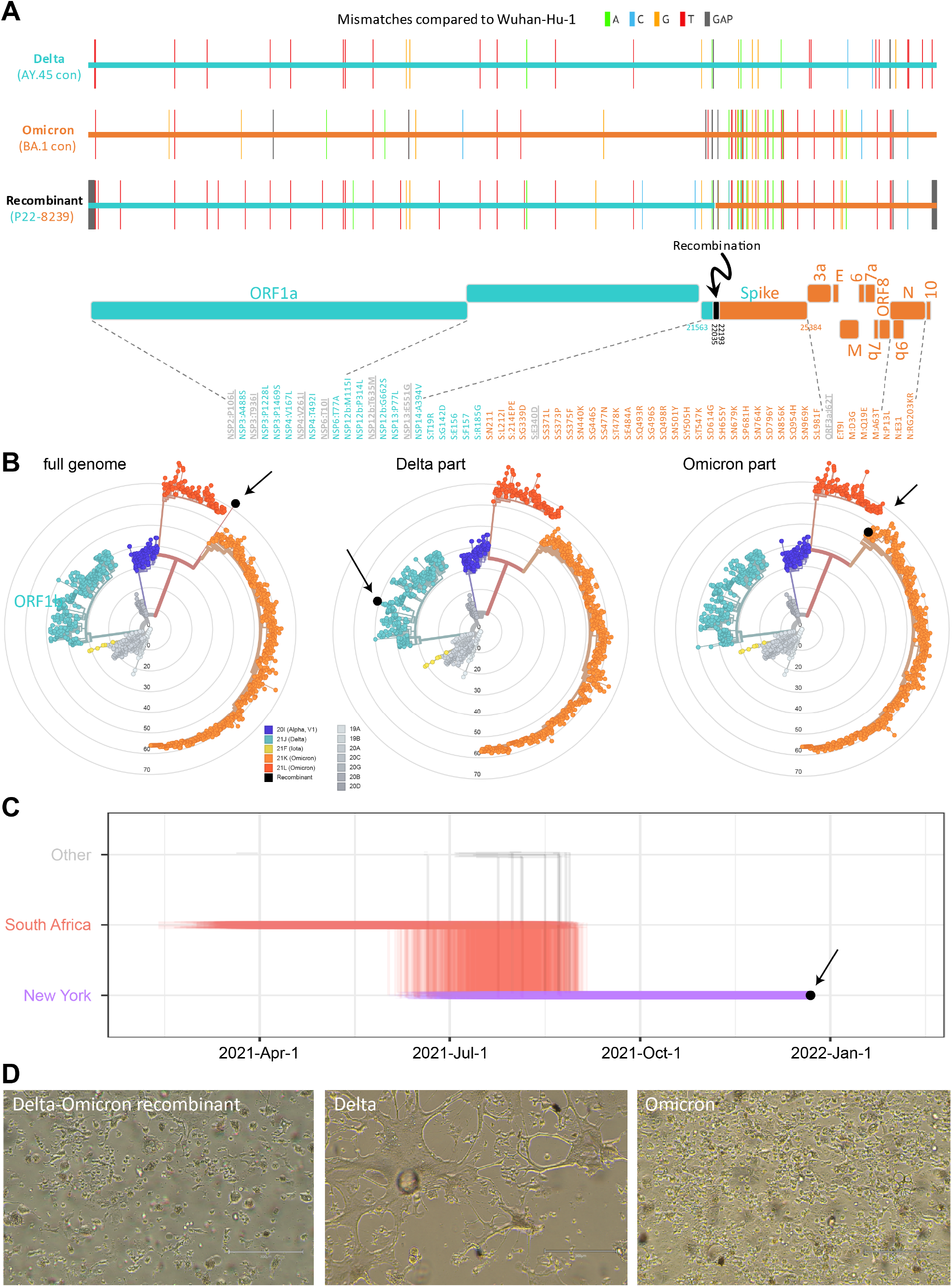
Genome mutations, phylogeographic analysis, and cytopathic effects of the Delta-Omicron recombinant SARS-CoV-2 virus. **A**. Full genome mutations of the SARS-CoV-2 Delta-Omicron recombinant, a Delta AY.45 consensus (con), and an Omicron BA.1 consensus sequence compared to the Wuhan-Hu-1 reference. Mutations are shown as ticks, color-coded according to the legend on the top. A schematic of the Delta and Omicron portions as well as the recombinant breakpoint region of the virus are shown. All non-synonymous mutations are listed and shown in teal (Delta mutation), orange (Omicron mutations), or gray underlined (not related to Delta or Omicron). **B**. Maximum likelihood phylogenies of 1557 global SARS-CoV-2 full genome sequences based on a New York, New Jersey, and Connecticut focused subsampling. Nextstrain clades are colored according to the legend and the recombinant variant is shown in black and highlighted by arrows. The phylogenetic analysis was done over the recombinant’s full genome (left) and separately for the Delta or Omicron parts only (middle and right), the latter by masking the complementary Omicron or Delta genomic regions, respectively. Concentrical rings indicate the number of mutations compared to the root Wuhan-Hu-1 sequence. **C**. Markov jump trajectory plot depicting the ancestral location history of the AY.45 segment of the recombinant sequence. Each individual trajectory corresponds to the Markov jumps in a single tree from the posterior distribution of 1,800 trees (**Figure S2B**). Horizontal lines represent the time maintained at an ancestral location and vertical lines represent a Markov jump between two locations. Despite the variability of topologies in the posterior trees, the trajectory plot for the AY.45 part of the recombinant genome consistently shows support for an ancestral history in South Africa, prior to infection of the patient in the state of New York as a result of local transmission (first SARS-CoV-2 positive test of patient is indicated in black and highlighted by an arrow). **D**. Cytopathic effects observed in various lineages of SARS-CoV-2 on VeroE6/TMPRSS2 cells. Omicron (BA.1, right) induces cell rounding and partial detachment from the monolayer, while Delta (AY.119, middle) induces large syncytia with extreme morphological changes and complete detachment. The Delta-Omicron recombinant (left) displayed cell rounding with eventual detachment from the monolayer, with large cells observed. Images were taken with an EVOS M5000 inverted microscope (ThermoFisher Scientific, Waltham, MA); 10X magnification.

### Phylogeographic origin of the Delta AY.45 infection lies in New York

While phylogenetic inference was not reliably feasible for BA.1 due to the large number of infections in conjunction with a low number of mutations, a maximum clade credibility (phylogenetic) tree using >1000 Delta AY.45 genomes and the Delta piece of the recombinant (**Figure S2**) revealed a monophyletic clade consisting entirely of United States AY.45 genomes including the recombinant (100% posterior support) as a descendant of a South African ancestral AY.45 backbone. The Markov jump trajectory plot (**Figure 1C**) backed an ancestral origin in South Africa and points to a jump from South Africa into New York state between June and July, 2021. The lineage associated with the recombinant genome was circulating in New York for >100 days before the positive COVID-19 test in December 2021, suggesting that infection with AY.45 occurred in New York. The phylogeographic reconstruction further shows that all US infections within this AY.45 clade originated from New York.

### Cytopathic effect of the recombinant is characteristic of Delta and Omicron

Cytopathic effects (CPE) of the *in vitro*-grown recombinant virus were first observed at 48 hpi on VeroE6/TMPRSS2 cells, with minor cell rounding in small areas of the monolayer, progressing to widespread cell rounding, detachment, debris, and cell death by 96 hpi (**Figure S1**), consistent with Omicron. However, affected cells appeared larger and more prominent in the CPE produced by the recombinant virus, and cell detachment was more advanced. As with Omicron, syncytia formation was not apparent in the recombinant virus cultures. Overall, the recombinant virus induced CPE more similar to those seen with Omicron than Delta (**Figure 1D**). By 96 hpi (Ct 13.5), viral production had increased more than 5 log^10^ from the 24h timepoint in the culture supernatant (**Table S2**).

### The recombinant breakpoint is adjacent to the quaternary Sotrovimab epitope in spike

Since recombination occurred in the carboxy-terminal portion of spike NTD (bp 22035-22193) (**Figure 1A**), most of the NTD is composed of Delta sequence, while RBD and the C-terminal regions are Omicron (**Figure 2**). During the first symptomatic COVID-19 episode in December 2021, the patient was treated with Sotrovimab, a class 3 anti-spike RBD nAb with broad antiviral activity against SARS-CoV-2, including Delta and BA.1^27-30^. The spike binding epitope for Sotrovimab involves the N-terminus of RBD, and, in a 3D structural model, the Fab moiety engages the space between RBD and the neighboring NTD (**Figure 2**). The recombinant virus sequence harbors one atypical spike mutation (E340D) not related to Delta or Omicron and rarely found in global SARS-CoV-2 sequences with a frequency of less than 3×10^−5 14,31^. Interestingly, E340 is in the middle of the Sotrovimab-binding epitope and is the primary site of resistance to Sotrovimab^27,32,33^. E340D is a conservative replacement without a change in charge, in contrast to the more abundant resistance mutations E340A/K^27,32^, yet we show that E340D can confer Sotrovimab resistance (**Figure 3**).

**Figure 2.**
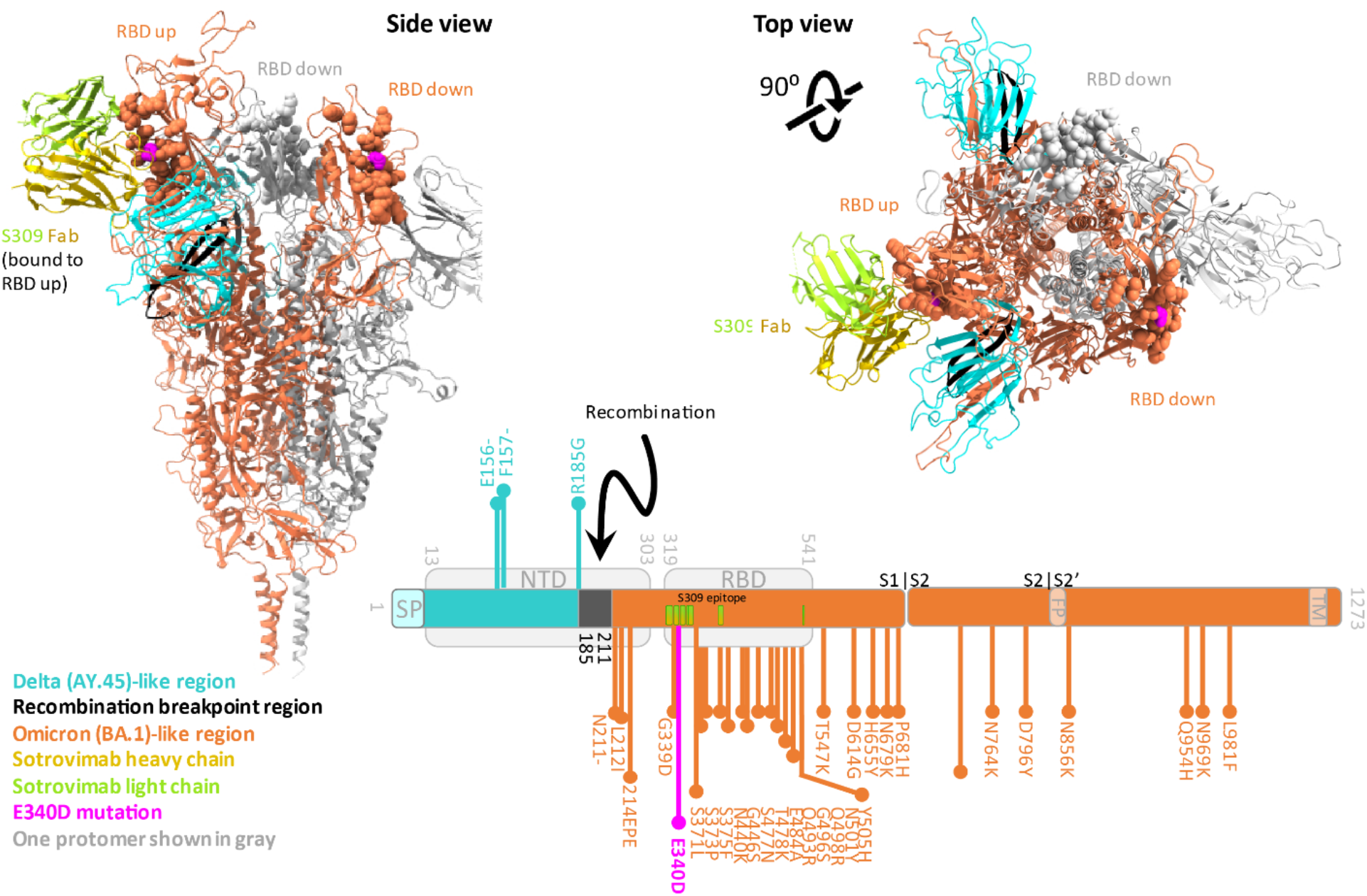
Breakpoint region of the Delta-Omicron recombinant spike in relation to the Sotrovimab (S309) binding epitope. Trimer spike structure of the Delta-Omicron recombinant in the open, one RBD-up conformation with one Sotrovimab Fab molecule bound to the RBD in up-position. Sotrovimab is a slightly refined, engineered version of its precursor S309 that has been used in the structures shown here. The spike portions are shown as ribbons and the Sotrovimab epitopes as spheres. One spike protomer is shown in gray; the other two protomers are colored according to the legend. Lower right: schematic of the location of the Delta (teal) and Omicron (orange) spike portions as well as the recombinant breakpoint region (black) including the mutations related to Delta (teal, facing up) or Omicron (orange, facing down). The mutation E340D, which is not related to Delta or Omicron, is highlighted in pink. The recombinant spike structure is a homology model of the recombinant spike based on pdb 7TO4 (1 RBD-up Omicron spike trimer). The S309 (Sotrovimab) molecule was added by structural overlay of a S309-bound spike co-structure in 1 RBD-up position (pdb 7TM0). FP: fusion peptide, NTD: N-terminal domain, RBD: receptor-binding domain, SP: signal peptide, TM: transmembrane domain.

**Figure 3.**
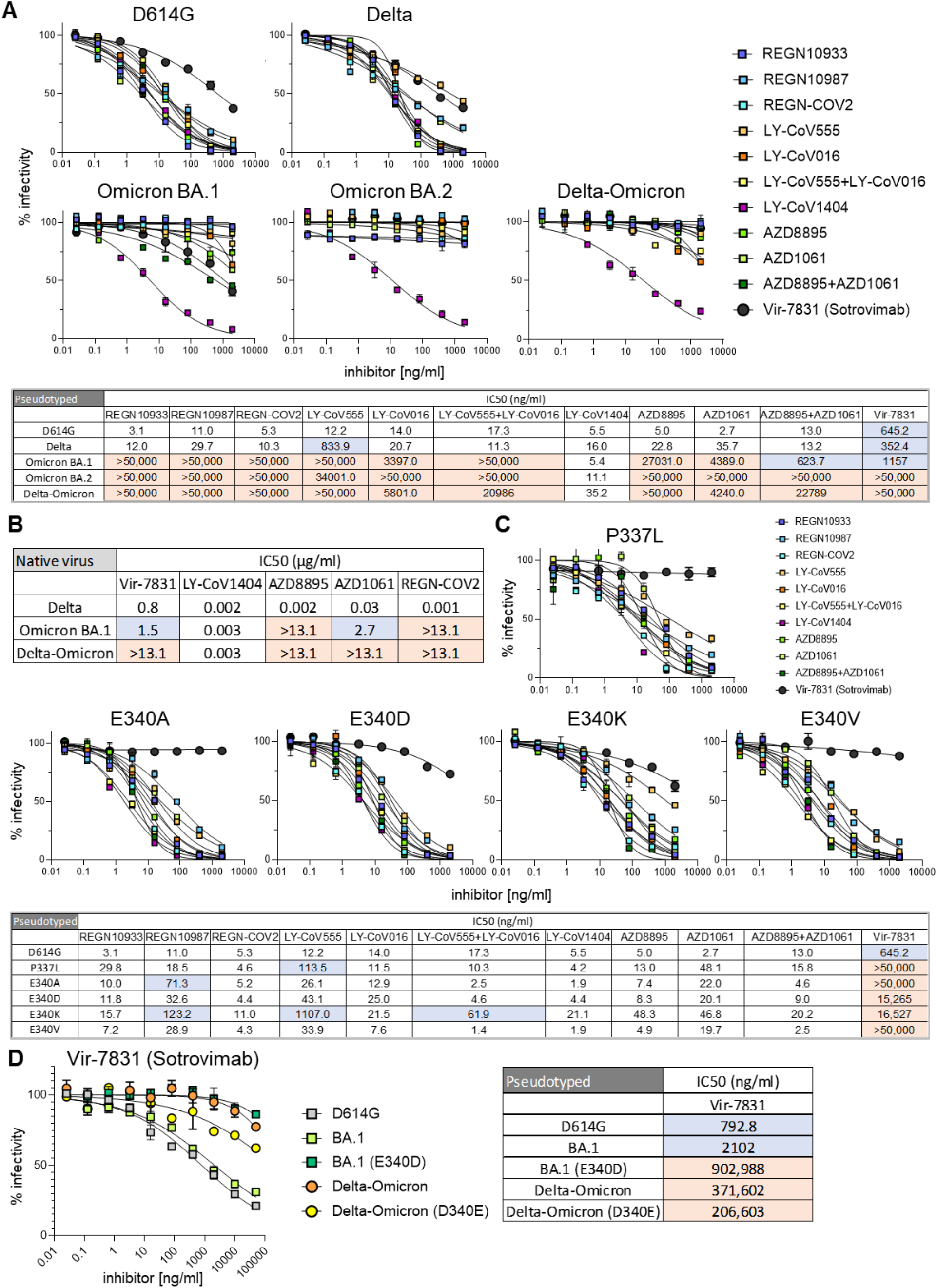
Delta-Omicron is resistant to most therapeutic monoclonal antibodies including Sotrovimab, irrespective of the E340D mutation in RBD. **A**. Neutralization of viruses with the D614G, Delta, Omicron, and Delta-Omicron spike proteins by the monoclonal antibodies (mAbs) Regeneron REGN10933 (Casirivamab), REGN10987 (Imdevimab), the REGN-COV-2 cocktail (Casirivimab+Imdevimab), LY-CoV555 (Bamlanivimab), LY-CoV016 (Etesevimab), LY-CoV1404 (Bebtelovimab), AZD8895 (Tixagevimab), AZD1061 (Cilgavimab), the AZD8895+AZD1061 combination (Evusheld), and VIR-7831 (Sotrovimab). The table shows the half-maximal inhibitory concentrations (IC50) of the therapeutic mAbs calculated using the data from the antibody neutralization curves shown above. Antibodies that did not allow the virus to reach >70% infectivity at the highest concentration tested are listed as IC50>50,000 ng/ml; for antibodies that reached 50-70% infectivity at the highest concentration tested, the IC50 was extrapolated. High resistance (IC50>3000 ng/ml) is highlighted in red, moderate resistance (50 ng/ml<IC50<3000 ng/ml) in blue, and no/low resistance (IC50<50 ng/ml) shown in white. **B**. IC50 values of therapeutic mAbs (selection of mAbs used in A) against infectious native virus calculated from plaque reduction neutralization tests (PNRTs, **Figure S3**). High resistance (IC50>10 μg/ml) is highlighted in red, moderate resistance (1 μg/ml<IC50<10 μg/ml) in blue, and no/low resistance (IC50<1 μg/ml) shown in white. **C**. Neutralization of D614G (backbone) and P337L, E340A, E340D, E340K and E340V point mutated spike protein-pseudotyped viruses by the same set of mAbs as in A. Experimental details and formatting as in A. **D**. Neutralization of D614G (control), BA.1 (E340), E340D point-mutated BA.1, Delta-Omicron (D340), and D340E point-mutated Delta-Omicron spike protein-pseudotyped viruses by VIR-7831 (Sotrovimab). Experimental details and formatting as in A.

### Delta-Omicron recombinant exhibits enhanced Sotrovimab resistance compared to Delta and BA.1

Delta viruses are sensitive to most therapeutic anti-SARS-CoV-2 antibodies, including all 11 mAbs and cocktails we tested, whereas BA.1 is resistant to most of them (8/11; **Figure 3A**). Among the few nAbs that retained activity against BA.1 are LY-CoV1404 and the cocktail AZD8895+AZD1061 (Evusheld) as well as the class 3 anti-RBD nAb VIR-7831 (Sotrovimab), the latter binding RBD outside the ACE2 binding site, in contrast to the more frequent class 1 and 2 nAbs^30^. Notably, while Delta and Omicron BA.1 are sensitive to Sotrovimab, the Delta-Omicron recombinant was resistant using pseudotyped (>43x increase in IC50 over BA.1) and infectious virus (>9x increase in IC50 over BA.1) (**Figures 3B, S3**). The Delta-Omicron recombinant was resistant to 10/11 tested nAbs, comparable to Omicron BA.2, and only the cocktail LY-CoV1404 retained activity (**Figure 3A**). Because the identified Delta-Omicron recombinant carries an unusual E340D mutation in the Sotrovimab epitope at a site that has been linked with Sotrovimab resistance (E340A/K)^32^, we compared the impact of E340D on Sotrovimab neutralization with other mutations that were recorded upon Sotrovimab treatment/resistance (**Figure 3C**). While E340A/V and P337L conferred complete Sotrovimab resistance in D614G pseudotyped virus (IC50>50,000 ng/ml), E340D’s impact was lower (IC50=15,265 ng/ml) but substantial (24x IC50 increase) and comparable to E340K (26x IC50 increase). All five mutations interfered with Sotrovimab neutralization exclusively but not with any of the other 10 nAbs tested (**Figure 3C**).

To deconvolute the mutual contributions of E340D and the recombination itself to Sotrovimab resistance, we performed site-directed mutagenesis in the recombinant and BA.1 pseudotyped viruses (**Figure 3D**). E340D in the BA.1 background conferred Sotrovimab resistance, as shown before for D614G. To test the consequence of Delta-Omicron recombination in the absence of E340D, we reverted 340D to 340E in the otherwise chimeric spike. Notably, the D340E revertant pseudotyped virus retained resistance to Sotrovimab (IC50>50,000 ng/ml) with only a subtle increase in sensitivity compared to the original (D340) version (1.8x change from 371,602 to 206,603 ng/ml). Thus, we conclude that recombination of Delta and Omicron spike was sufficient to confer resistance to Sotrovimab, and that E340D slightly enhanced the resistance phenotype (**Figure 3D**).

### Delta-Omicron virus resistance against polyclonal antibody responses and clinical impact of E340X and P337L mutations

The Delta-Omicron recombinant was resistant to naturally and vaccine-induced anti-SARS-CoV-2 nAbs to a level comparable with that of BA.1 or BA.2 (**Figure 4A, Table S3**). The neutralization sensitivity of the recombinant remained low (IC50<600 serum dilution) even in COVID-19-experienced individuals 1 month after boost, compared to IC50s>10,000 with Delta or D614G viruses. In boosted individuals, the recombinant had 2.5x (COVID-19-naïve) and 1.9x (COVID-19-experienced) lower IC50 values compared to BA.1, suggesting a slightly, though non-significantly enhanced resistant phenotype. Immune responses induced 1 month post 2^nd^ vaccination were significantly reduced against D614G virus carrying E340A/V/D or P337L (but not E340K) (**Figure 4B**). This finding suggests that Sotrovimab-like RBD class 3 nAbs are present in vaccinee sera and that Sotrovimab resistance mutations have an impact on vaccine-induced polyclonal nAb responses.

**Figure 4.**
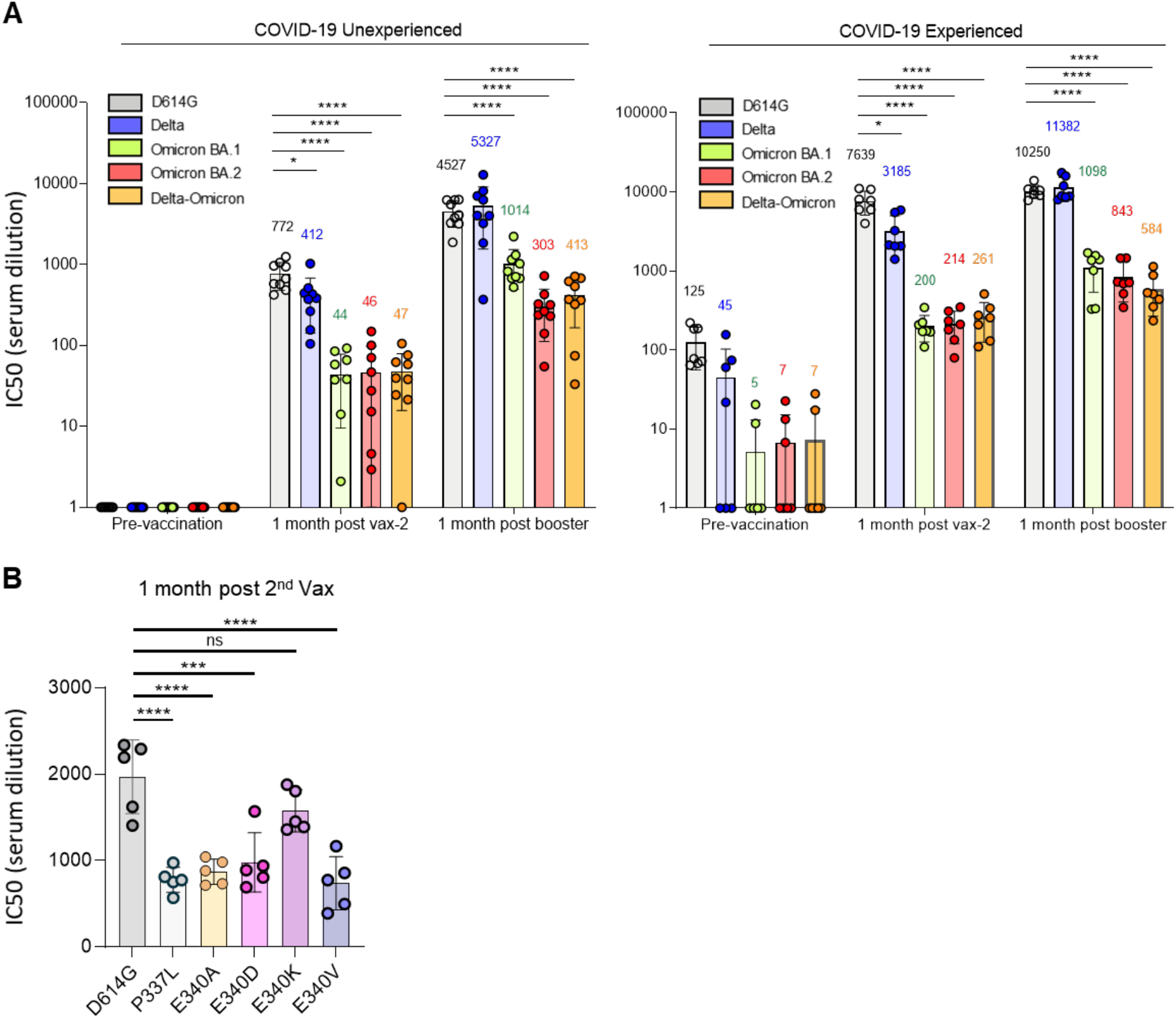
Resistance of the Delta-Omicron recombinant to vaccine-elicited antibodies. **A**. Neutralizing antibody titers of participants with or without previous history of SARS-CoV-2 infection and/or vaccination were measured using pseudotyped viruses with D614G, Delta, Omicron, and Delta-Omicron spike. Sera were collected from study participants pre-vaccination, 1-month post-second vaccination with Pfizer BNT162b2, and 1-month post-boost. Study participants were without previous SARS-CoV-2 infection (unexperienced) (left) (n=9) or previously infected (experienced) (right) (n=7). COVID-19 history was determined by symptoms and a PCR+ test or serology. Equivalent amounts of D614G, Delta, Omicron, and Delta-Omicron spike protein-pseudotyped viruses were mixed with a 2-fold serial dilution of donor serum and then applied to ACE2.293T cells. Luciferase activity was measured two days post-infection. Each serum dilution was measured in triplicate and the experiment was done twice with similar results. Half-maximal inhibitory concentrations (IC50) are shown for one representative experiment. Mean values for each group are shown above the bar. **B**. Neutralizing antibody titers of sera from COVID-19 unexperienced individuals (n=5) collected 1 month after 2^nd^ vaccination against D614G (as backbone) and P337L, E340A, E340D, E340K and E340V point mutated spike protein-pseudotyped viruses. Statistical significance was calculated by two-sided testing. (*P≤0.05, ***P≤0.001, ****P≤0.0001).

## Discussion

Drivers for recombinant variant selection have remained obscure, and functionally distinctive features of SARS-CoV-2 recombinants are scarcely known. In contrast to other recombinants reported so far^6,8,12,34^, we describe a SARS-CoV-2 recombinant detected in an immunosuppressed transplant recipient after mAb treatment (**Figure 1A,B**). Immunocompromised patients are a primary source of new variants due to prolonged viral replication/evolution, particularly under selective pressure by mAb treatment^35-38^, which, in the case we describe, may have resulted in a mAb-resistant recombinant variant. We show for the first time that recombination between Delta and BA.1, two variants intrinsically sensitive to Sotrovimab, rendered the recombinant spike resistant to Sotrovimab (**Figure 3**). These results highlight recombination as a mechanism for SARS-CoV-2 neutralization escape.

We showed that the Delta-Omicron recombinant evaded neutralization by Sotrovimab, as well as other mAbs and vaccine-elicited antibodies (**Figures 3**,**4**). In line with the latter finding, a recent study suggested a more vaccine-resistant phenotype of a Delta-Omicron pseudotyped virus compared with D614G and BA.3^16^; mAbs were not tested. Another report using a spike reporter assay indicated that Delta-Omicron recombinants have similar resistance as BA.1 to natural infection or COVID-19 vaccine-induced nAbs but less pronounced as for BA.4/5^17^.

Among the reported recombinant SARS-CoV-2 sequences, others, and ours show a single breakpoint splitting the genome into a 5’ Delta and a 3’ Omicron segment^6,8^. A three-case cluster in France was caused by a recombinant with a smaller Omicron portion, principally spike, flanked by 5’ and 3’ Delta regions^12^. All these cases share an ORF1ab Delta region and an Omicron region encompassing spike’s RBD and C-terminal regions, suggesting that either a recombination breakpoint in spike NTD is favored, or Delta’s non-structural proteins or Omicron’s C-terminal spike regions, including RBD, have selective advantages.

Notably, we detected the recombinant virus in a patient after treatment with Sotrovimab, a nAb binding SARS-CoV-2 spike near the recombination site. High-resolution structures will be needed to determine whether recombination altered the conformation/accessibility of the Sotrovimab epitope, thus rendering the virus resistant to Sotrovimab. Our neutralization data further showed that both the recombination event in spike NTD and the RBD mutation E340D added up to a combined resistance phenotype (**Figure 3D**). It is plausible that Sotrovimab treatment selected for the recombinant and/or its E340D mutation, and recombination and E340D co-evolved reinforcing each other. Since Delta had disappeared from circulation by February 2022 in the NYC metro area, and presuming that in December 2021 Sotrovimab selected for the E340D mutation in the Omicron part of spike, we speculate that the patient became co-infected with Delta and Omicron in December 2021 when Delta and Omicron were at ∿10% and 90% prevalence in New York, respectively. Alternatively, the participant became infected with the recombinant from an undefined source. Our titration experiments suggest that E340D confers partial Sotrovimab escape comparable to E340K when tested against D614G background virus. E340D, located in the center of the Sotrovimab binding epitope in RBD, is extremely rare and neither related to a specific lineage nor community spread, and might thus be an indicator of exerted Sotrovimab immune pressure. Besides mAbs (Sotrovimab), E340D confers resistance to polyclonal Abs (**Figures 3C**,**4B**) and thus needs to be considered an immune escape mutation.

Since the recombinant was the only variant detected in the February 2022 specimen without parental Delta or Omicron lineages present, we can only speculate whether the recombinant was generated after Delta/Omicron co-infection in December 2021 with subsequent escape from and selection against Sotrovimab. Phylogenetically related BA.1 and AY.45 sequences (**Figure 1C**) were highly prevalent in New York at this time, which render a regional Delta/Omicron coinfection and recombination event more likely compared to the introduction of an unknown recombinant from an undefined source. Delta-Omicron recombinants remained rare without rampant community transmissions, possibly because of their selective advantage against monoclonal but less against polycloncal Abs (**Figures 3A**,**4A**), thus primarily affecting high-risk/immunocompromised patients treated with therapeutic mAbs. This case stresses the importance of genomic surveillance and functional testing to identify new mechanisms and pathways of immune escape, e.g., by novel recombinant forms. Monitoring of SARS-CoV-2 infections in immunocompromised individuals treated with mAbs remains a priority for pandemic preparedness to detect neutralization escape variants.

## Acknowledgements

We thank Dr. Joan Cangiarella for support of SARS-CoV-2 genomic surveillance at NYULH and for institutional funding. We thank Drs Marie I. Samanovic-Golden and Mark J. Mulligan from the NYU Langone Vaccine Center and the Department of Medicine for the clinical samples and related information. We are grateful to the authors and submitting laboratories who deposited data in GISAID, in particular to those whose sequences we used to create the phylogenetic tree. We thank Cornelius Roemer from the Biozentrum, University of Basel and Nextclade for initial assessment of our sequence. The project was partially supported by the NYU Clinical and Translational Science award 3UL1TR001445-06A1S1, and the NYULH Genome Technology Center is partially supported by the Cancer Center Support Grant P30CA016087 at the Laura and Isaac Perlmutter Cancer Center. The work was further funded by grants from the NIH to N.R.L. (DA046100, AI122390 and AI120898). T.T. was supported by the Vilcek/Goldfarb Fellowship Endowment Fund. SLH and GB acknowledge support from the Research Foundation - Flanders (”Fonds voor Wetenschappelijk Onderzoek - Vlaanderen,” G0E1420N). GB acknowledges support from the Internal Funds KU Leuven (Grant No. C14/18/094) and the Research Foundation - Flanders (”Fonds voor Wetenschappelijk Onderzoek - Vlaanderen,” G098321N, V420922N). We also thank the staff of the Wadsworth Center Applied Genomics Technology Core for performing ARTIC sequencing. The sequencing by the Wadsworth Center is supported in part by Cooperative Agreement Number NU50CK000516, funded by the Centers for Disease Control and Prevention. The contents of this paper are solely the responsibility of the authors and do not necessarily represent the official views of the Centers for Disease Control and Prevention or the Department of Health and Human Services.

## References

1. Simon-Loriere E, Holmes EC. Why do RNA viruses recombine? Nature Reviews Microbiology 2011;9(8):617–626. DOI: 10.1038/nrmicro2614.

2. Bean WJ, Jr., Cox NJ, Kendal AP. Recombination of human influenza A viruses in nature. Nature 1980;284(5757):638-40. (In eng). DOI: 10.1038/284638a0.

3. Moradigaravand D, Kouyos R, Hinkley T, et al. Recombination Accelerates Adaptation on a Large-Scale Empirical Fitness Landscape in HIV-1. PLOS Genetics 2014;10(6):e1004439. DOI: 10.1371/journal.pgen.1004439.

4. Zhu Z, Meng K, Meng G. Genomic recombination events may reveal the evolution of coronavirus and the origin of SARS-CoV-2. Scientific Reports 2020;10(1):21617. DOI: 10.1038/s41598-020-78703-6.

5. Pollett S, Conte MA, Sanborn M, et al. A comparative recombination analysis of human coronaviruses and implications for the SARS-CoV-2 pandemic. Scientific Reports 2021;11(1):17365. DOI: 10.1038/s41598-021-96626-8.

6. Bolze A, White S, Basler T, et al. Evidence for SARS-CoV-2 Delta and Omicron coinfections and recombination. medRxiv 2022:2022.03.09.22272113. DOI: 10.1101/2022.03.09.22272113.

7. Rockett RJ, Draper J, Gall M, et al. Co-infection with SARS-CoV-2 Omicron and Delta variants revealed by genomic surveillance. Nature Communications 2022;13(1):2745. DOI: 10.1038/s41467-022-30518-x.

8. Lacek KA, Rambo-Martin BL, Batra D, et al. Identification of a Novel SARS-CoV-2 Delta-Omicron Recombinant Virus in the United States. bioRxiv 2022:2022.03.19.484981. DOI: 10.1101/2022.03.19.484981.

9. Jackson B, Boni MF, Bull MJ, et al. Generation and transmission of interlineage recombinants in the SARS-CoV-2 pandemic. Cell 2021;184(20):5179-5188.e8. DOI: 10.1016/j.cell.2021.08.014.

10. Sekizuka T, Saito M, Itokawa K, et al. Recombination between SARS-CoV-2 Omicron BA.1 and BA.2 variants identified in a traveller from Nepal at the airport quarantine facility in Japan. Journal of Travel Medicine 2022. DOI: 10.1093/jtm/taac051.

11. Gu H, Ng DYM, Liu GYZ, et al. Recombinant BA.1/BA.2 SARS-CoV-2 Virus in Arriving Travelers, Hong Kong, February 2022. Emerg Infect Dis 2022;28(6):1276-1278. (In eng). DOI: 10.3201/eid2806.220523.

12. Colson P, Fournier PE, Delerce J, et al. Culture and identification of a “Deltamicron” SARS-CoV-2 in a three cases cluster in southern France. J Med Virol 2022;94(8):3739-3749. (In eng). DOI: 10.1002/jmv.27789.

13. Wang L, Zhou HY, Li JY, et al. Potential Inter-variant and Intra-variant Recombination of Delta and Omicron Variants. J Med Virol 2022 (In eng). DOI: 10.1002/jmv.27939.

14. Hadfield J, Megill C, Bell SM, et al. Nextstrain: real-time tracking of pathogen evolution. Bioinformatics 2018;34(23):4121–4123. DOI: 10.1093/bioinformatics/bty407.

15. Duerr R, Dimartino D, Marier C, et al. Clinical and genomic signatures of rising SARS-CoV-2 Delta breakthrough infections in New York. medRxiv 2021:2021.12.07.21267431. DOI: 10.1101/2021.12.07.21267431.

16. Evans JP, Qu P, Zeng C, et al. Neutralization of the SARS-CoV-2 Deltacron and BA.3 Variants. New England Journal of Medicine 2022;386(24):2340–2342. DOI: 10.1056/NEJMc2205019.

17. Kurhade C, Zou J, Xia H, et al. Neutralization of Omicron sublineages and Deltacron SARS-CoV-2 by 3 doses of BNT162b2 vaccine or BA.1 infection. bioRxiv 2022:2022.06.05.494889. DOI: 10.1101/2022.06.05.494889.

18. Duerr R, Dimartino D, Marier C, et al. Dominance of Alpha and Iota variants in SARS-CoV-2 vaccine breakthrough infections in New York City. J Clin Invest 2021;131(18). DOI: 10.1172/JCI152702.

19. Matsuyama S, Nao N, Shirato K, et al. Enhanced isolation of SARS-CoV-2 by TMPRSS2-expressing cells. Proc Natl Acad Sci U S A 2020;117(13):7001–7003. DOI: 10.1073/pnas.2002589117.

20. Los Alamos National Laboratory tools. Highlighter tool. (http://www.hiv.lanl.gov/).

21. Suchard MA, Lemey P, Baele G, Ayres DL, Drummond AJ, Rambaut A. Bayesian phylogenetic and phylodynamic data integration using BEAST 1.10. Virus Evol 2018;4(1):vey016. DOI: 10.1093/ve/vey016.

22. Goddard TD, Huang CC, Meng EC, et al. UCSF ChimeraX: Meeting modern challenges in visualization and analysis. Protein Sci 2018;27(1):14-25. (In eng). DOI: 10.1002/pro.3235.

23. Waterhouse A, Bertoni M, Bienert S, et al. SWISS-MODEL: homology modelling of protein structures and complexes. Nucleic Acids Res 2018;46(W1):W296-w303. (In eng). DOI: 10.1093/nar/gky427.

24. Tada T, Zhou H, Dcosta BM, et al. Increased resistance of SARS-CoV-2 Omicron variant to neutralization by vaccine-elicited and therapeutic antibodies. EBioMedicine 2022;78:103944. DOI: 10.1016/j.ebiom.2022.103944.

25. Tada T, Fan C, Chen JS, et al. An ACE2 Microbody Containing a Single Immunoglobulin Fc Domain Is a Potent Inhibitor of SARS-CoV-2. Cell Rep 2020;33(12):108528. DOI: 10.1016/j.celrep.2020.108528.

26. O’Toole Á, Scher E, Underwood A, et al. Assignment of epidemiological lineages in an emerging pandemic using the pangolin tool. Virus Evolution 2021;7(2). DOI: 10.1093/ve/veab064.

27. Rockett R, Basile K, Maddocks S, et al. Resistance Mutations in SARS-CoV-2 Delta Variant after Sotrovimab Use. New England Journal of Medicine 2022. DOI: 10.1056/NEJMc2120219.

28. Gupta A, Gonzalez-Rojas Y, Juarez E, et al. Early Treatment for Covid-19 with SARS-CoV-2 Neutralizing Antibody Sotrovimab. New England Journal of Medicine 2021;385(21):1941–1950. DOI: 10.1056/NEJMoa2107934.

29. Cameroni E, Bowen JE, Rosen LE, et al. Broadly neutralizing antibodies overcome SARS-CoV-2 Omicron antigenic shift. Nature 2022;602(7898):664–670. DOI: 10.1038/s41586-021-04386-2.

30. Barnes CO, Jette CA, Abernathy ME, et al. SARS-CoV-2 neutralizing antibody structures inform therapeutic strategies. Nature 2020;588(7839):682–687. DOI: 10.1038/s41586-020-2852-1.

31. CoV-GLUE. CoV-GLUE - Amino acid replacements. (http://cov-glue.cvr.gla.ac.uk/#/replacement).

32. Starr TN, Czudnochowski N, Liu Z, et al. SARS-CoV-2 RBD antibodies that maximize breadth and resistance to escape. Nature 2021;597(7874):97–102. DOI: 10.1038/s41586-021-03807-6.

33. Birnie E, Biemond JJ, Appelman B, et al. Development of Resistance-Associated Mutations After Sotrovimab Administration in High-risk Individuals Infected With the SARS-CoV-2 Omicron Variant. JAMA 2022. DOI: 10.1001/jama.2022.13854.

34. Cheng Y. Possible English recombinant descended from Omicron and Delta (UKHSA “signal under monitoring”) #422. (https://github.com/cov-lineages/pango-designation/issues/422).

35. Choi B, Choudhary MC, Regan J, et al. Persistence and Evolution of SARS-CoV-2 in an Immunocompromised Host. New England Journal of Medicine 2020;383(23):2291–2293. DOI: 10.1056/NEJMc2031364.

36. Weigang S, Fuchs J, Zimmer G, et al. Within-host evolution of SARS-CoV-2 in an immunosuppressed COVID-19 patient as a source of immune escape variants. Nature Communications 2021;12(1):6405. DOI: 10.1038/s41467-021-26602-3.

37. Nussenblatt V, Roder AE, Das S, et al. Yearlong COVID-19 Infection Reveals Within-Host Evolution of SARS-CoV-2 in a Patient With B-Cell Depletion. J Infect Dis 2022;225(7):1118-1123. (In eng). DOI: 10.1093/infdis/jiab622.

38. Kemp SA, Collier DA, Datir RP, et al. SARS-CoV-2 evolution during treatment of chronic infection. Nature 2021;592(7853):277–282. DOI: 10.1038/s41586-021-03291-y.

