## Supplemental Appendix for "Delta-Omicron recombinant escapes therapeutic antibody neutralization"

This appendix has been provided by the authors to give readers additional information about their work.

#### List of investigators

Ralf Duerr, MD[1,*,#], Hao Zhou, PhD[1,#], Takuya Tada, PhD[1,#], Dacia Dimartino, PhD[2], Christian Marier, BSc[2], Paul Zappile, MS[2], Guiqing Wang, MD[3], Jonathan Plitnick, MA[4], Sara B. Griesemer, MS[4], Roxanne Girardin, PhD[4], Erica Lasek-Nesselquist, PhD[5,6], Samuel L. Hong, BA[7], Guy Baele, PhD[7], Meike Dittmann, PhD[1], Mila B. Ortigoza, MD[1,6], Prithiv J. Prasad, MD[6], Kathleen McDonough, PhD[4,8], Nathaniel R. Landau, PhD[1], Kirsten St. George, PhD[4,8], and Adriana Heguy, PhD [2,3,*]
* Co-corresponding authors
### Shared contribution

Correspondence:
Ralf Duerr, MD, PhD
Department of Microbiology, NYU Grossman School of Medicine
Alexandria Center for Life Science (ACLS), West Tower
430 East, 29th Street, Room 323
New York, NY 10016

&
Adriana Heguy, PhD
Department of Pathology, NYU Grossman School of Medicine
Genome Technology Center, NYU Langone Health
550 First Avenue, MSB 294A
New York, NY 10018


#### Affiliations

1. Department of Microbiology, NYU Grossman School of Medicine
2. Genome Technology Center, Office of Science and Research, NYU Langone Health
3. Department of Pathology, NYU Grossman School of Medicine
4. Laboratory of Viral Diseases, Wadsworth Center, New York State Department of Health, Albany, NY
5. Bioinformatics Core, Wadsworth Center, New York State Department of Health, Albany, NY
6. Department of Medicine, NYU Grossman School of Medicine
7. Department of Microbiology, Immunology and Transplantation, Laboratory for Clinical and Epidemiological Virology, Rega Institute, KU Leuven, Leuven, Belgium
8. Biomedical Sciences Department, School of Public Health, University at Albany, SUNY, Albany, NY

#### Supplemental Methods

**SARS-CoV-2 sequencing and bioinformatic analysis**

1. IDT xGen Amplicon and metagenomics approach (NYU).

Genomic surveillance was carried out as described previously [1], using an amplicon-based library prep method. The web-based Nextclade v1.14 [2] and Auspice v2.36 phylogenomic visualization features of Nextstrain [3] were used to examine viral genome clade assignment and mutation calling. For metagenomics, a shotgun library from total RNA extracted from the nasal swab was generated using the Illumina Stranded Total RNA prep Ligation with Ribo-Zero Plus. Sequencing was performed on an Illumina Nova Seq 6000, 150PE, dual index.

Sequencing reads were demultiplexed using the Illumina bcl2fastq2 Conversion Software v2.20 and adapters and low-quality bases were trimmed with Trimmomatic v0.36 [4]. BWA v0.7.17 [5] was utilized for mapping reads to the SARS-CoV-2 reference genome (NC_045512.2, wuhCor1) and duplicate reads were removed using Sambamba v0.6.8 [6]. GATK v3.8 DepthOfCoverage and HaplotypeCaller tools [7] were used to determine on-target viral coverage and call mutations.

2. Ion AmpliSeq Insight and ARTIC amplicon-based approach (NY State DOH).

Total nucleic acid was extracted from the primary clinical specimen and samples of cultured isolate using the bioMerieux NucliSens® easyMAG® platform (bioMerieux Inc, Durham, NC), using 110 uL of sample. Real-time RT-PCR was performed on the original primary specimen, as well as the cultured isolate, using the New York SARS-CoV-2 Real-Time Reverse Transcriptase (RT)-PCR Diagnostic Panel, according to the protocol indicated in the Instructions for Use [8]. Library preparation was performed on RNA from primary specimen with an Ion Chef and sequencing with an Ion S5 XL, using the Ion AmpliSeq SARS-CoV-2 Insight Research Assay (ThermoFisher Scientific, Waltham, MA, USA). The assay was performed with 27 cycles and a 4-minute extension time. Library preparation was performed on RNA from the cultured isolates using a modified

ARTIC protocol with ARTIC V4 primers, as previously described [9]. Libraries were sequenced on an Illumina NextSeq instrument.

Consensus genomes were generated as previously described[9] with the ncov2019-artic-nf pipeline (https://github.com/connor-lab/ncov2019-artic-nf). Mutations were identified with Nextclade (https://clades.nextstrain.org/) and lineage assignment was performed with Pangolin v3.1.19 (pangoLearn 2022-01-22)[10] for both consensus genomes, which identified the first part of the recombinant genome as AY.45, i.e., a sublineage of the Delta variant of concern (VOC), and the second part as Omicron BA.1. Additionally, we used Usher (https://genome.ucsc.edu/cgi-bin/hgPhyloPlace ) to place consensus genomes in a global SARS-CoV-2 phylogenetic tree to identify closest relatives. Lineage composition of the sequencing libraries was determined by Freyja (https://github.com/andersen-lab/Freyja) to evaluate the possibility of a mixed infection.

**Cells and media**

VeroE6/TMPRSS2 cells, a modified VeroE6 cell line expressing the transmembrane serine protease, TMPRSS2 [11], were used for virus isolation, obtained from the Japanese Collection of Research Bioresources (JCRB Cell Bank) cell number JCRB1819, through Sekisui Xenotech, LLC (Kansas City, KS), agreement # A2000230. Cells were maintained in Dulbecco's modified Eagle's medium (DMEM) supplemented with sodium bicarbonate and 10% fetal bovine serum (FBS, all from Millipore Sigma, St. Louis, MO), as well as 1mg/1mL geneticin G418 (Gibco).

**Virus isolation**

VeroE6/TMPRSS2 cells were seeded in T25 flasks with 5mL cell suspension (1.5 x $10^5$ cells/mL), three days prior to infection, to reach 85-90% confluency. Before inoculation, 300 μL of specimen were combined with 300 μL penicillin/streptomycin (pen/strep, 10 units/mL Millipore Sigma, St. Louis, MO), 150 μL of nystatin (1mg/mL; Millipore Sigma), and 250 μL of gelatin-Tris-Hanks (GTH). The cell monolayer was infected with the processed specimen, adsorbed 1.5h at 37$^o$C with 5% $CO_2$, then overlaid with 5 mL viral growth medium as described above, except FBS was reduced to 2%. In accordance with NY State DOH safety policy, the SARS-CoV-2 inoculations were performed at biosafety level 2 (BSL-2) but all subsequent incubations, culture samplings and harvesting were performed at biosafety level 3 (BSL-3). Infected monolayers were checked daily for cytopathic effect (CPE), and 110 μL of supernatant were removed at 24, 48, 72, and 96 hours post-infection (hpi) and immediately placed in NucliSENS lysis buffer for further analysis. At 96 hpi, cells and supernatant were harvested together, and 110uL of the harvest was also lysed for further analysis. All samples were extracted and tested by real-time RT-PCR to confirm viral growth.

**Mutation analysis**

Highlighter analyses were performed on MAFFT-aligned SARS-CoV-2 full-genome sequences using the Highlighter tool provided by the Los Alamos HIV sequence database [12] against Wuhan-Hu-1 as master. Delta AY.45 and Omicron BA.1 consensus sequences were used for comparison. They were generated with the EMBOSS cons tool (50% consensus threshold) [13] using complete, high-coverage AY.45 and BA.1 sequences from North-America sampled between June and December 2021 and downloaded from GISAID [14].

**Phylogenetic analysis**

Phylogenetic analyses were done using the Nextstrain CLI package WSL on Windows [3]. As input, we used the full genome sequence of the recombinant virus (hCoV-19/USA/NY-NYULH6045/2022) and a North-America-focused global dataset until the end of February 2022 (n=2259), complemented with a global Delta AY.45 data set of complete, high-coverage sequences from November 2021 until the end of February 2022 (n=335), all of which were downloaded (FASTA files and metadata) from GISAID [14]. The trees were constructed using a focal-contextual subsampling, focusing on sequences from the Tri-state area of New York, New Jersey, and Connecticut (including the recombinant) and complemented with a maximum of 1500 contextual sequences outside the Tri-state area and prioritized by proximity to the focal data set (total of n=1557 sequences). Default parameters were used including the masking of the first 100 and last 50 bp of the SARS-CoV-2 alignment and creating a maximum-likelihood IQ time-calibrated tree, rooted to Wuhan-Hu-1[3]. The phylogenetic analyses of the Delta and Omicron subregions of the recombinant were done by masking the complementary regions (Omicron or Delta genomic regions, respectively) and skipping the minimum length criterium.

**Phylogeographic analysis**

We performed Bayesian phylogeographic analysis to determine the plausible geographic origins of AY.45 and BA.1 infections. We combined the AY.45 part of the recombinant with all available high-quality (with high coverage, low coverage excluded and with complete sampling dates) AY.45 genomes on GISAID [14] until December 31st, 2021. We used Nextalign v1.11.0 [2] to align all genomes to the reference and trimmed the resulting alignment. Next, we estimated an unrooted maximum-likelihood phylogeny using IQ-TREE v2.2.0 [15] with automated model selection, which determined the GTR+F+R3 model as yielding the highest fit to the data. We performed 1,000 bootstrap replicates using the ultrafast bootstrap approximation (UFBOOT) in IQ-TREE v2.2.0. Given the high bootstrap support obtained for the clade that contains the recombinant (bootstrap >= 92%), we performed a subsequent discrete phylogeographic analysis [16] with Bayesian stochastic search variable selection (BSSVS) using BEAST v1.10.5 [17] in combination with BEAGLE v3.2 [18] on a large subclade of the consensus phylogeny that consists of 1,122 AY.45 genomes, by employing an empirical tree distribution with default priors. In order to attain proper statistical mixing, we grouped USA states that only reported a single AY.45 genome into a "USA other" location state and removed all other locations with only a single AY.45 genome. We inspected convergence and mixing aspects of all relevant parameters using Tracer 1.7 [19] to ensure that their associated effective sample size (ESS) values were all >200. After having discarded 10% of sampled posterior trees as burn-in, we constructed a maximum clade credibility (MCC) tree using TreeAnnotator 1.10.5[17].

Given the large number of available BA.1 genomes until December 31st, 2021, the analysis pipeline as described for AY.45 could not be performed. We hence used the BA.1 part of the recombinant genome and employed BLAST to consider only the closest sequences to the BA.1 recombinant segment. We first downloaded and aligned all BA.1 genomes collected up to December 31st, 2021 (n=138,350) from GISAID and removed identical sequences originating from the same country or state in the case of sequences originating from the US. We used the resulting alignment of 105,176 sequences to create a local BLAST database, against which we queried the BA.1 part of the recombinant genome with a megablast search in BLAST+ v2.12.0 [20]. We allowed for a minimum word size of 100 and a maximum number of 10 high scoring segment pairs per target sequence in order to obtain as many non-overlapping local alignments. After manually checking that segments do not overlap, we aggregated all hits on the same target sequence and sorted by highest combined bitscore. A total of 4,706 sequences originating from 79 different countries were tied with the highest combined bitscore. We subsequently proceeded to align these highest scoring BA.1 genomes and obtain an unrooted maximum-likelihood phylogeny as described for the AY.45 data set, with IQ-TREE v2.2.0 this time selecting the GTR+F+R2 substitution model as the optimal model choice. However, the bootstrap analysis yielded an almost entirely unresolved phylogeny in the form of a large multifurcation (data not shown). As a result, no subsequent phylogeographic analysis was performed.

**Structural analysis**

Molecular graphics and analyses were performed with UCSF ChimeraX 1.3 [21]. Homology models of the recombinant spike were generated with the protein structure homology-modelling server SWISS-MODEL[22]. The template structure was chosen based on coverage, sequence similarity, Global Model Quality Estimation (GMQE) and Quaternary Structure Quality Estimation (QSQE) scores, oligo state, and open/closed state (pdb 7TO4, 1 RBD-up Omicron spike trimer). The Fab moiety of S309 (Sotrovimab) was added to the recombinant spike trimer model by structural overlay of a S309-bound spike co-structure in the open (1 RBD-up) position (pdb 7TM0) using MatchMaker with pairing of the RBDs in up position.

**Infectious virus neutralization assay**

Wadsworth Center plaque reduction neutralization tests (PRNTs) were performed by mixing 100 µl of low passage, sequence-confirmed SARS-CoV-2 virus isolates containing approximately 100-180 plaque forming units with 100 µl of 2-fold serially diluted monoclonal antibody preparations or test sera and incubating at 37°C in 5% $CO_2$ for one hour. Confluent Vero E6 with TMPRSS2 cells (JCRB1819, Sekusi XeonTech) seeded in 6 well plates were inoculated with 100 µl of the virus: antibody mixtures and adsorption proceeded for one hour at 37°C in 5% $CO_2$. A 0.6% agar overlay prepared in maintenance medium (Dulbecco's Modified Eagle Medium, 2% heat-inactivated fetal bovine serum, 100 µg/ml Penicillin G, 100 U/ml Streptomycin, 1mg/ml Geneticin) was added after adsorption and the assay was incubated at 37°C in 5% $CO_2$. A second agar overlay containing 0.2% Neutral red added was added two days post infection. The number of plaques in each well was recorded after two additional days of incubation. All procedures were conducted in biosafety level 3 (BSL-3) laboratory conditions. Viruses used include BA.1: hCoV-19/USA/NY-Wadsworth-21103366-01/2021; Delta: hCoV-19/USA/NY-Wadsworth-2200014356-01/2021; Delta-Omicron. $IC_{50}$ values were calculated by determining the percent neutralization (relative to virus only controls) for technical duplicates from two to four biological replicates of each well and using non-linear regression ([inhibitor] vs. normalized response with variable slope, GraphPad Prism).

**Pseudotyped virus neutralization assay**

*Plasmids*

The SARS-CoV-2 BA.1-spike was previously generated[23]. The Delta-like N-terminal domain of the SARS-CoV-2 Delta-Omicron recombinant spike was chemically synthesized as a short fragment (Genscript Biotech Corporation, Piscataway, New Jersey, USA). The N-terminal fragment was amplified with a forward primer containing a Kpn-I site and a reverse primer containing an inner Omicron BA.1 sequence. This fragment was fused with another fragment which encodes RBD and C-terminal parts of Omicron BA.1 spike including an Xho-I site by overlapping PCR. 19 amino acids at the spike C-terminus, a reported endoplasmic reticulum retention sequence, were deleted (Δ19)[24]. The full Delta-Omicron recombinant spike was cloned into the Kpn-I and Xho-I sites of pcDNA6 (Invitrogen). Point mutations were introduced by overlap extension. HIV-1 Gag/Pol expression vector pMDL and HIV-1 Rev expression vector pRSV.Rev have been previously described[25].

*Human sera and monoclonal antibodies*

Sera from individuals vaccinated at NYULH with BNT162b2 were collected through the NYU Vaccine Center with written consent under I.R.B. approval and were deidentified.

*Cells*

293T cells were cultured in Dulbecco's modified Eagle medium (DMEM) supplemented with 10% fetal bovine serum (FBS) and penicillin/streptomycin (P/S) at 37°C in 5% $CO_2$. Stable ACE2.293T cell line were cultured in DMEM supplemented with 10% fetal bovine serum (FBS) and penicillin/streptomycin (P/S) and 1 μg/ml puromycin, as previously described[25].

*SARS-CoV-2 spike lentiviral pseudotypes and neutralization assay*

SARS-CoV-2 variant spike pseudotyped lentivirus stocks were produced by cotransfection of 293T cells with pMDL, pLenti.GFP-NLuc, pcCoV2.S-Δ19 (or variants thereof) and pRSV.Rev as previously described[25]. Real-time PCR reverse transcriptase activity was used to measure the lentivirus stocks and normalize them[26]. To measure neutralizing titer, sera or mAbs were serially diluted (2-fold for sera and 5-fold for mAbs) and incubated for 30 min at room temperature with pseudotyped virus. The mixture was added on 1 X $10^4$ of ACE2.293T cells (MOI of 0.2). After 1 day of infection, medium was removed and infectivity was measured with Nano-Glo luciferase substrate (Nanolight). Luminescence was read in an Envision 2103 microplate luminometer (PerkinElmer). D614G virus (original Wuhan virus carrying the D614G spike mutation that was selected early during the pandemic) was used as reference.

*Quantification and Statistical Analysis*

All experiments were performed in technical duplicates and data were analyzed using GraphPad Prism 8. Statistical significance was determined by two-tailed, unpaired t-tests or one-way ANOVA (*$P \leq 0.05$, **$P \leq 0.01$, ***$P \leq 0.001$, ****$P \leq 0.0001$). Confidence intervals are shown as the mean ± standard deviation.

**Data sharing statement**
The sequence of the recombinant virus is publicly available in GISAID (hCoV-19/USA/NY-NYULH6045/2022; EPI_ISL_10792641).

#### Supplemental Results

**Multimethod SARS-CoV-2 sequencing confirms the identity of the recombinant variant**
The full-genome SARS-CoV-2 sequence was initially determined by xGen amplicon sequencing of the nasopharyngeal swab. The original swab was then re-extracted, re-sequenced, and processed in the same manner, which revealed the same 5' Delta and 3' Omicron-specific mutations with a breakpoint between 22,035-22,193 bp in the near full-length high-quality sequence (**Table S1**). No mixed bases were observed, suggesting that this was not a co-infection or technical artifact. For further confirmation, we prepared and sequenced libraries from the same sample using a ribodepletion shotgun metagenomics approach. GATK variant detection revealed a 100% concordance between mutations called with both the amplicon and independent shotgun metagenomics approach. The metagenomics approach yielded a single additional Omicron-specific change at the beginning of the Omicron-like portion of the genome, i.e., a 9 bp insertion at position 22,204 (spike 214EPE). The specimen was also separately extracted and sequenced by AmpliSeq Insight at a second site, NY State Department of Health (DOH) at Wadsworth, further confirming the sequencing results. Additionally, the virus was grown (NY State DOH) in VeroE6/TMPRSS2 cells and the progeny sequenced using another amplicon-based method, ARTIC V4, producing a fourth confirmation of the recombinant identity. Delta and Omicron-derived mutations were supported by 99-100% of the reads, consistent with the presence of a single recombinant. ARTIC sequencing confirmed the spike 214EPE insertion, but did not cover three spike mutations (K417N, N440K, G446S), detected by the previous approaches. **Table S1** summarizes all mutations detected by nine multi-method sequencing runs on different specimens.

**Phylogeographic origin of the recombinant's Delta AY.45 piece.**
We focused our deeper phylogenetic and phylogeographic analysis on the AY.45 part of the recombinant genome, since AY.45 had not been reported in other Delta-Omicron recombinants[27-29]. BA.1 was the dominant lineage at this time, leading to a large number of infections in conjunction with a low number of mutations, rendering reliable phylogenetic inference infeasible for the BA.1 part[30]. In a maximum clade credibility (phylogenetic) tree linking 1122 AY.45 genomes (**Figure S2A**), the AY.45 partial genome of the recombinant was located on a distinct branch (highlighted). A discrete phylogeographic analysis of the entire data set shows the ancestral location state reconstruction on a monophyletic cluster (100% posterior support) that contains the recombinant genome (**Figure S2B**). A monophyletic clade consisting entirely of US AY.45 genomes (100% posterior support) can be seen to descend from a South African ancestral AY.45 backbone, offering support for a South African origin for this particular clade of US AY.45 infections. The Markov jump trajectory plot[31] for the Delta part of the recombinant (**Figure 1C**) also shows strong support for an ancestral origin in South Africa. Our phylogeographic

reconstruction points to a jump from South Africa into the state of New York in the period from mid-June until the end of July, 2021. This reconstruction shows that the lineage associated with the recombinant genome was circulating in New York for 115 to 184 days before the patient testing COVID-19 positive in late December 2021, suggesting that infection with AY.45 occurred in the state of New York. The phylogeographic reconstruction further shows that all infections in the US that fall within this AY.45 clade originated from New York state.

#### References

1. Duerr R, Dimartino D, Marier C, et al. Dominance of Alpha and Iota variants in SARS-CoV-2 vaccine breakthrough infections in New York City. J Clin Invest 2021;131(18). DOI: 10.1172/JCI152702.
2. Aksamentov I, Roemer C, Hodcroft EB, Neher RA. Nextclade: clade assignment, mutation calling and quality control for viral genomes (1.4.5). Zenodo 2021 (https://doi.org/10.5281/zenodo.5726681).
3. Hadfield J, Megill C, Bell SM, et al. Nextstrain: real-time tracking of pathogen evolution. Bioinformatics 2018;34(23):4121-4123. DOI: 10.1093/bioinformatics/bty407.
4. Bolger AM, Lohse M, Usadel B. Trimmomatic: a flexible trimmer for Illumina sequence data. Bioinformatics 2014;30(15):2114-20. DOI: 10.1093/bioinformatics/btu170.
5. Li H, Durbin R. Fast and accurate short read alignment with Burrows-Wheeler transform. Bioinformatics 2009;25(14):1754-60. DOI: 10.1093/bioinformatics/btp324.
6. Tarasov A, Vilella AJ, Cuppen E, Nijman IJ, Prins P. Sambamba: fast processing of NGS alignment formats. Bioinformatics 2015;31(12):2032-4. DOI: 10.1093/bioinformatics/btv098.
7. McKenna A, Hanna M, Banks E, et al. The Genome Analysis Toolkit: a MapReduce framework for analyzing next-generation DNA sequencing data. Genome Res 2010;20(9):1297-303. (In eng). DOI: 10.1101/gr.107524.110.
8. Wadsworth Center NYSDoH. New York SARS-CoV-2 Real-Time Reverse Transcriptase (RT)-PCR Diagnostic Panel, Instructions for Use. . (https://www.fda.gov/media/135847/download).
9. Plitnick J, Griesemer S, Lasek-Nesselquist E, Singh N, Lamson DM, St George K. Whole-Genome Sequencing of SARS-CoV-2: Assessment of the Ion Torrent AmpliSeq Panel and Comparison with the Illumina MiSeq ARTIC Protocol. J Clin Microbiol 2021;59(12):e0064921. DOI: 10.1128/JCM.00649-21.
10. O'Toole Á, Scher E, Underwood A, et al. Assignment of epidemiological lineages in an emerging pandemic using the pangolin tool. Virus Evolution 2021;7(2). DOI: 10.1093/ve/veab064.
11. Matsuyama S, Nao N, Shirato K, et al. Enhanced isolation of SARS-CoV-2 by TMPRSS2-expressing cells. Proc Natl Acad Sci U S A 2020;117(13):7001-7003. DOI: 10.1073/pnas.2002589117.
12. Los Alamos National Laboratory tools. Highlighter tool. (http://www.hiv.lanl.gov/).
13. explorer; E. EMBOSS cons. (https://www.bioinformatics.nl/cgi-bin/emboss/cons).
14. Shu Y, McCauley J. GISAID: Global initiative on sharing all influenza data - from vision to reality. Euro Surveill 2017;22(13) (In eng). DOI: 10.2807/1560-7917.Es.2017.22.13.30494.
15. Minh BQ, Schmidt HA, Chernomor O, et al. IQ-TREE 2: New Models and Efficient Methods for Phylogenetic Inference in the Genomic Era. Mol Biol Evol 2020;37(5):1530-1534. (In eng). DOI: 10.1093/molbev/msaa015.
16. Lemey P, Rambaut A, Drummond AJ, Suchard MA. Bayesian phylogeography finds its roots. PLoS Comput Biol 2009;5(9):e1000520. DOI: 10.1371/journal.pcbi.1000520.

17. Suchard MA, Lemey P, Baele G, Ayres DL, Drummond AJ, Rambaut A. Bayesian phylogenetic and phylodynamic data integration using BEAST 1.10. Virus Evol 2018;4(1):vey016. DOI: 10.1093/ve/vey016.
18. Ayres DL, Cummings MP, Baele G, et al. BEAGLE 3: Improved Performance, Scaling, and Usability for a High-Performance Computing Library for Statistical Phylogenetics. Systematic Biology 2019;68(6):1052-1061. DOI: 10.1093/sysbio/syz020.
19. Rambaut A, Drummond AJ, Xie D, Baele G, Suchard MA. Posterior Summarization in Bayesian Phylogenetics Using Tracer 1.7. Syst Biol 2018;67(5):901-904. DOI: 10.1093/sysbio/syy032.
20. Morgulis A, Coulouris G, Raytselis Y, Madden TL, Agarwala R, Schäffer AA. Database indexing for production MegaBLAST searches. Bioinformatics 2008;24(16):1757-64. (In eng). DOI: 10.1093/bioinformatics/btn322.
21. Goddard TD, Huang CC, Meng EC, et al. UCSF ChimeraX: Meeting modern challenges in visualization and analysis. Protein Sci 2018;27(1):14-25. (In eng). DOI: 10.1002/pro.3235.
22. Waterhouse A, Bertoni M, Bienert S, et al. SWISS-MODEL: homology modelling of protein structures and complexes. Nucleic Acids Res 2018;46(W1):W296-w303. (In eng). DOI: 10.1093/nar/gky427.
23. Tada T, Zhou H, Dcosta BM, et al. Increased resistance of SARS-CoV-2 Omicron variant to neutralization by vaccine-elicited and therapeutic antibodies. EBioMedicine 2022;78:103944. DOI: 10.1016/j.ebiom.2022.103944.
24. Giroglou T, Cinatl J, Rabenau H, et al. Retroviral Vectors Pseudotyped with Severe Acute Respiratory Syndrome Coronavirus S Protein. Journal of Virology 2004;78(17):9007-9015. DOI: doi:10.1128/JVI.78.17.9007-9015.2004.
25. Tada T, Fan C, Chen JS, et al. An ACE2 Microbody Containing a Single Immunoglobulin Fc Domain Is a Potent Inhibitor of SARS-CoV-2. Cell Rep 2020;33(12):108528. DOI: 10.1016/j.celrep.2020.108528.
26. Vermeire J, Naessens E, Vanderstraeten H, et al. Quantification of Reverse Transcriptase Activity by Real-Time PCR as a Fast and Accurate Method for Titration of HIV, Lenti- and Retroviral Vectors. PLOS ONE 2012;7(12):e50859. DOI: 10.1371/journal.pone.0050859.
27. Lacek KA, Rambo-Martin BL, Batra D, et al. Identification of a Novel SARS-CoV-2 Delta-Omicron Recombinant Virus in the United States. bioRxiv 2022:2022.03.19.484981. DOI: 10.1101/2022.03.19.484981.
28. Bolze A, White S, Basler T, et al. Evidence for SARS-CoV-2 Delta and Omicron co-infections and recombination. medRxiv 2022:2022.03.09.22272113. DOI: 10.1101/2022.03.09.22272113.
29. Colson P, Fournier PE, Delerce J, et al. Culture and identification of a "Deltamicron" SARS-CoV-2 in a three cases cluster in southern France. J Med Virol 2022;94(8):3739-3749. (In eng). DOI: 10.1002/jmv.27789.
30. Morel B, Barbera P, Czech L, et al. Phylogenetic Analysis of SARS-CoV-2 Data Is Difficult. Molecular Biology and Evolution 2020;38(5):1777-1791. DOI: 10.1093/molbev/msaa314.
31. Lemey P, Hong SL, Hill V, et al. Accommodating individual travel history and unsampled diversity in Bayesian phylogeographic inference of SARS-CoV-2. Nature Communications 2020;11(1):5110. DOI: 10.1038/s41467-020-18877-9.

#### Supplemental Figures

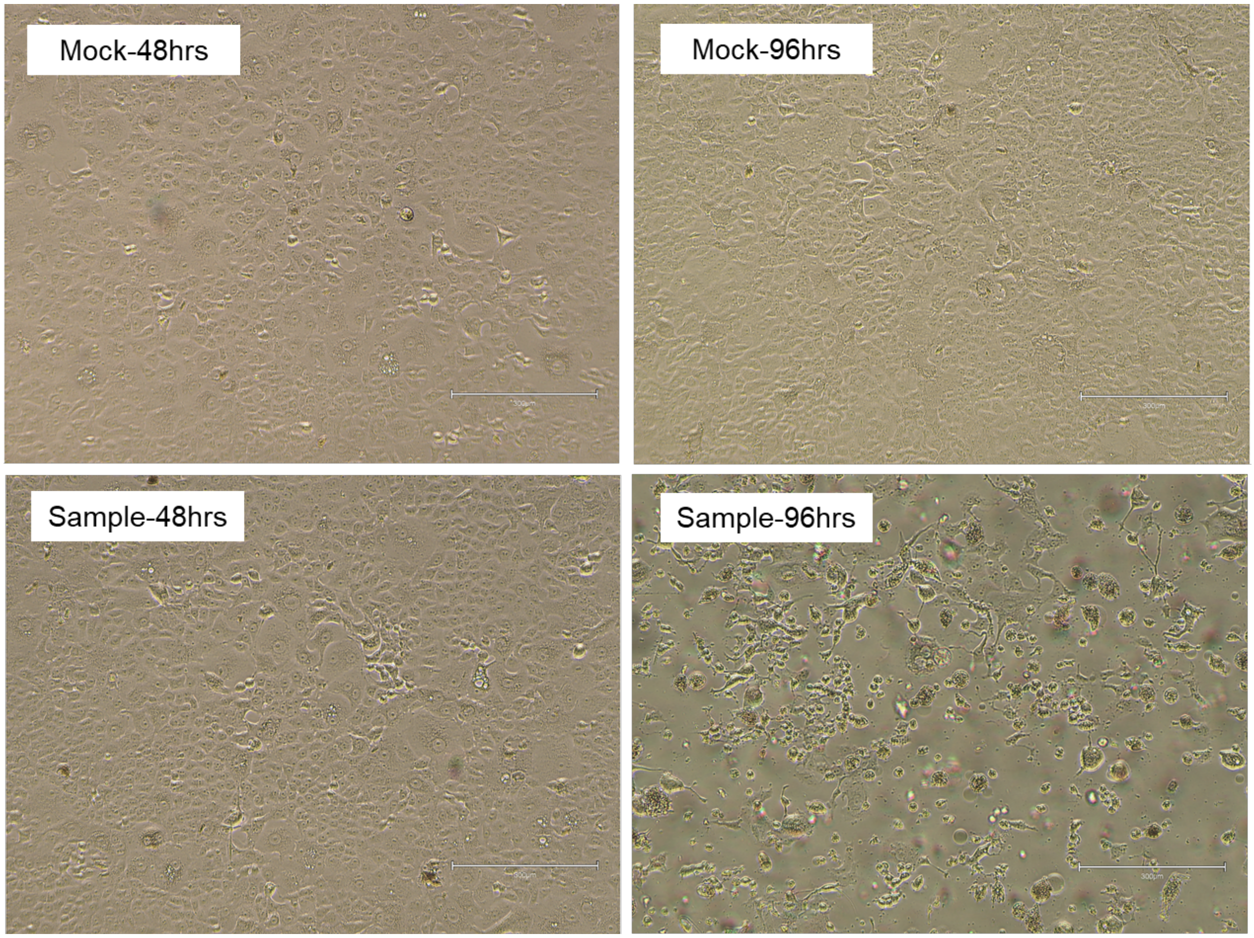

**Figure S1. Cytopathic effect of the SARS-CoV-2 Delta-Omicron recombinant on VeroE6/TMPRSS2 cells.**

A residual nasopharyngeal swab specimen of the patient infected with the recombinant variant was inoculated onto VeroE6/TMPRSS2 cells and observed daily for cytopathic effects (bottom row). The image on the lower right (Sample-96hrs) is the same as the left image in **Figure 1D** (Delta-Omicron recombinant). Images were taken with an EVOS M5000 inverted microscope (ThermoFisher Scientific, Waltham, MA); 10X magnification.

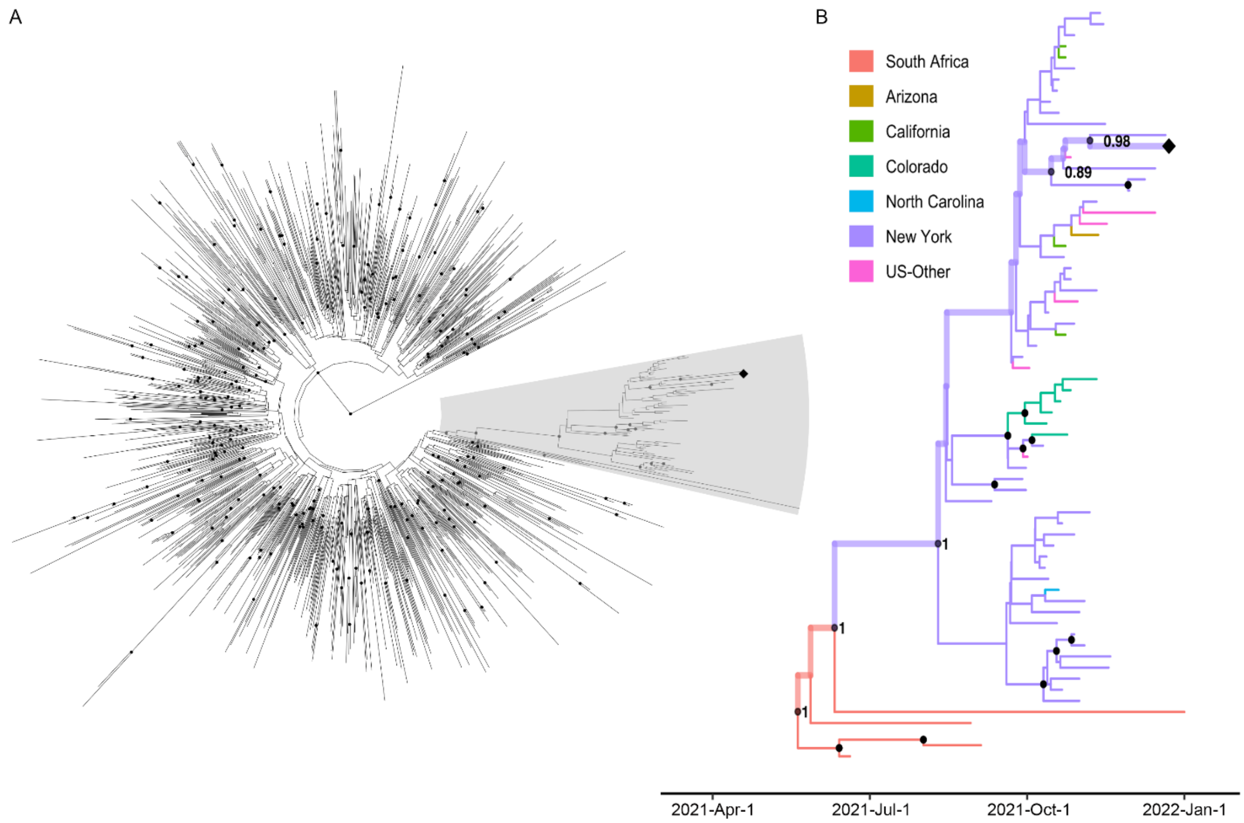

**Figure S2. Phylogeographic analysis of the AY.45 cluster containing the recombinant.**
**A.** Maximum clade credibility tree of 1122 AY.45 genomes. The recombinant genome (AY.45 segment of it) is highlighted with a diamond symbol. Nodes with a posterior support above 70% are denoted with a black circle. A well supported phylogenetic cluster containing the recombinant genome is shaded in gray. **B.** Phylogeographic analysis of the phylogenetic cluster containing the AY.45 segment of the recombinant sequence. The recombinant is annotated with a diamond symbol, and branches corresponding to the Markov jump trajectory plot (**Figure 1C**) are highlighted (by thicker branches). Internal nodes with posterior support of 70% or higher are denoted with a black circle, whereas nodes with 100% support are annotated explicitly.

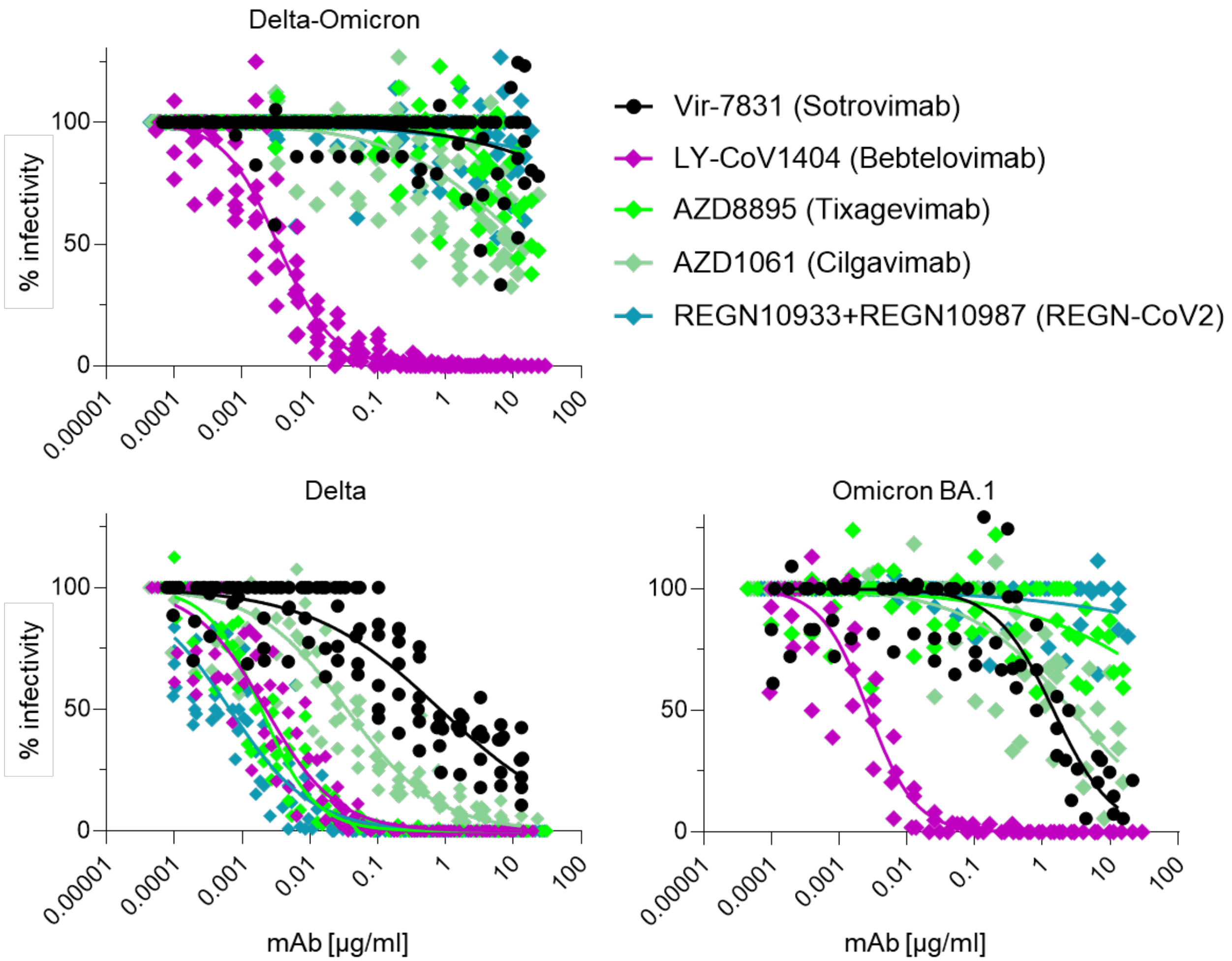

**Figure S3. Native Delta-Omicron virus is resistant to several therapeutic monoclonal antibodies including Sotrovimab.**

Neutralization of infectious Delta-Omicron virus in comparison with Delta and BA.1 by five different monoclonal antibodies (mAbs) including Vir-7831 (Sotrovimab), LY-CoV1404 (Bebtelovimab), AZD8895 (Tixagevimab), AZD1061 (Cilgavimab), and REGN-COV2 (Casirivimab & Imdevimab). Inhibition was determined in plaque reduction neutralization tests using serially diluted mAb doses. The graphs display all data points (staggered) from two to four biological replicates with technical duplicates (Delta-Omicron: four biological replicates). Neutralization curves are shown as non-linear regression fits for each mAb.

#### Supplemental Tables

**Table S1. Summary of mutations detected in the Delta-Omicron recombinant by four sequencing methods on different sample aliquots, with or without *in vitro* culture.**

| gene | nucleotide | amino acid | donor | xGen | xGen (repeat) | shotgun | AmpliSeq | harvest | 24h | 48h | 72h | 96h |
|---|---|---|---|---|---|---|---|---|---|---|---|---|
| ORF1ab | C1122T | P106L | Delta | X | X | X | X | X | X | X | X | X |
| ORF1ab | G4181T | A488S | Delta | X | X | X | X | X | X | X | X | X |
| ORF1ab | C5526T | T936I | Delta | X | X | X | X | X | X | X | X | X |
| ORF1ab | C6402T | P1228L | Delta | X | X | X | X | X | X | X | X | X |
| ORF1ab | C7124T | P1469S | Delta | X | X | X | X | X | X | X | X | X |
| ORF1ab | G9053T | V167L | Delta | X | X | X | X | X | X | X | X | X |
| ORF1ab | G9335A | V261I | Delta | X | X | X | X | X | X | X | X | X |
| ORF1ab | C10029T | T492I | Delta | X | X | X | X | X | X | X | X | X |
| ORF1ab | C11001T | T10I | Delta | X | X | X | X | X | X | X | X | X |
| ORF1ab | A11201G | T77A | Delta | X | X | X | X | X | X | X | X | X |
| ORF1ab | G13812T | M124I | Delta | X | X | X | X | X | X | X | X | X |
| ORF1ab | C14408T | P323L | Delta | X | X | X | X | X | X | X | X | X |
| ORF1ab | C15371T | T644M | Delta | X | X | X | X | X | X | X | X | X |
| ORF1ab | G15451A | G671S | Delta | X | X | X | X | X | X | X | X | X |
| ORF1ab | C16466T | P77L | Delta | X | X | X | X | X | X | X | X | X |
| ORF1ab | A17888G | E551G | Delta | X | X | X | X | X | X | X | X | X |
| ORF1ab | C19220T | A394V | Delta | X | X | X | X | X | X | X | X | X |
| Spike | C21618G | T19R | Delta | X | X | X | X | X | X | X | X | X |
| Spike | G21987A | G142D | Delta | X | X | X | X | X | X | X | X | X |
| Spike | GAGTTCA22028G | EFR156-158G deletion | Delta | X | X | X | X | X | X | X | X | X |
| Spike | AATT22193A | NL211-212I deletion | Omicron | X | X | X | X | X | X | X | X | X |
| Spike | T22204TGAGCCAGAA | 214EPE insertion | Omicron | -- | -- | X | X | X | X | X | X | X |
| Spike | G22578A | G339D | Omicron | X | X | X | X | X | X | X | X | X |
| Spike | A22582C | E340D | Omicron | X | X | X | X | X | X | X | X | X |
| Spike | T22673C | S371P | Omicron | X | X | X | X | X | X | X | X | X |
| Spike | C22674T | S371F | Omicron | X | X | X | X | X | X | X | X | X |
| Spike | T22679C | S373P | Omicron | X | X | X | X | X | X | X | X | X |
| Spike | C22686T | S375F | Omicron | X | X | X | X | X | X | X | X | X |
| Spike | G22813T | K417N | Omicron | X | X | X | X | -- | -- | -- | -- | -- |
| Spike | T22882G | N440K | Omicron | X | X | X | X | -- | -- | -- | -- | -- |
| Spike | G22898A | G446S | Omicron | X | X | X | X | -- | -- | -- | -- | -- |
| Spike | G22992A | S477N | Omicron | X | X | X | X | X | X | X | X | X |
| Spike | C22995A | T478K | Omicron | X | X | X | X | X | X | X | X | X |
| Spike | A23013C | E484A | Omicron | X | X | X | X | X | X | X | X | X |
| Spike | A23040G | Q493R | Omicron | X | X | X | X | X | X | X | X | X |
| Spike | G23048A | G496S | Omicron | X | X | X | X | X | X | X | X | X |

| | | | | | | | | | | | | |
|---|---|---|---|---|---|---|---|---|---|---|---|---|
| Spike | A23055G | Q498R | Omicron | X | X | X | -- | X | X | X | X | X |
| Spike | A23063T | N501Y | Omicron | X | X | X | -- | X | X | X | X | X |
| Spike | T23075C | Y505H | Omicron | X | X | X | -- | X | X | X | X | X |
| Spike | C23202A | T547K | Omicron | X | X | X | X | X | X | X | X | X |
| Spike | A23403G | D614G | Omicron | X | X | X | X | X | X | X | X | X |
| Spike | C23525T | H655Y | Omicron | X | X | X | X | X | X | X | X | X |
| Spike | T23599G | N679K | Omicron | X | X | X | X | X | X | X | X | X |
| Spike | C23604A | P681H | Omicron | X | X | X | X | X | X | X | X | X |
| Spike | C23854A | N764K | Omicron | X | X | X | X | X | -- | X | X | X |
| Spike | G23948T | D796Y | Omicron | X | X | X | X | X | X | X | X | X |
| Spike | C24130A | N856K | Omicron | X | X | X | X | X | X | X | X | X |
| Spike | A24424T | Q954H | Omicron | X | X | X | X | X | X | X | X | X |
| Spike | T24469A | N969K | Omicron | X | X | X | X | X | X | X | X | X |
| Spike | C24503T | L981F | Omicron | X | X | X | X | X | X | X | X | X |
| ORF3a | T25577C | I62T | Omicron | X | X | X | X | X | X | X | X | X |
| E | C26270T | T9I | Omicron | X | X | X | X | X | X | X | X | X |
| M | A26530G | D3G | Omicron | X | X | X | X | X | X | X | X | X |
| M | C26577G | Q19E | Omicron | X | X | X | X | X | X | X | X | X |
| M | G26709A | A63T | Omicron | X | X | X | X | X | X | X | X | X |
| N | C28311T | P13L | Omicron | X | X | X | X | X | X | X | X | X |
| N | GGAGAACGCA28361G | ERS31-33 deletion | Omicron | X | X | X | X | X | X | X | X | X |
| N | G28881A | R203K | Omicron | X | X | X | X | X | X | X | X | X |
| N | G28883C | G204R | Omicron | X | X | X | X | X | X | X | X | X |

xGen: IDT xGen NGS amplicon sequencing (performed at NYU);
shotgun: metagenomics shotgun sequencing (performed at NYU;
AmpliSeq: AmpliSeq Insight sequencing (performed at NY State DOH);
harvest: ARTIC NGS amplicon sequencing using cells and supernatant after 96h virus culture;
24h-96h: ARTIC NGS amplicon sequencing using supernatants after the indicated culture time points (performed at NY State DOH);
--: no coverage

**Table S2. Real-time RT-PCR results during culture of the SARS-CoV-2 Delta-Omicron recombinant in VeroE6/TMPRSS2 cells.**

| Sample | Ct value (N1) |
|---|---|
| NPS specimen | 30.8 |
| 24hpi | 30.6 |
| 48hpi | 23.1 |
| 72hpi | 16.8 |
| 96hpi | 13.5 |

Abbreviations: Ct: cycle threshold value; hpi: hours post infection; NPS: naso-pharyngeal swab.

**Table S3. IC50 of BNT162b2-elicited antibodies against viruses with variant spike proteins in sera collected from COVID-19-unexperienced and experienced donors (related to Figure 4A,B)**

| BNT162b2, COVID-19-Unexperienced | | | | | | | | | | | |
|---|---|---|---|---|---|---|---|---|---|---|---|
| | | | | D614G | | Delta | | Omicron BA.1 | | Omicron BA.2 | |
| Donor | Age | Sex | Comorbidities | 1 month post vax-2 | 1 month post booster | 1 month post vax-2 | 1 month post booster | 1 month post vax-2 | 1 month post booster | 1 month post vax-2 | 1 month post booster |
| 1 | 34 | M | None | 1250 | 4106 | 105 | 371 | 37.84 | 1027 | 15.27 | 326.5 |
| 2 | 29 | F | None | 572 | 3206 | 156.2 | 1851 | 14.11 | 824.8 | 4.605 | 238.9 |
| 3 | 37 | F | Allergy | 946 | 6359 | 386 | 3234 | 58.34 | 677.5 | 70.71 | 319.1 |
| 4 | 62 | M | Hypertension, Hyperlipidemia | 671.2 | 3789 | 1032 | 4024 | 85 | 1154 | ND | 428.8 |
| 5 | 52 | F | Hypertesion | 581.1 | 3275 | 265.4 | 7040 | 66.11 | 2221 | 149.1 | 141 |
| 6 | 34 | M | Hypothyroidism | 966.2 | 6301 | 433.1 | 6319 | ND | 527 | 100.6 | 54.87 |
| 7 | 38 | F | Asthma, Anemia, Tinea versicolor | 1060 | 5541 | 444.8 | 4045 | 2.097 | 1315 | ND | 724.6 |
| 8 | 38 | M | None | 423.3 | 1882 | 373.2 | 8163 | 38.33 | 678.7 | 27.7 | 231.6 |
| 9 | 52 | F | None | 484.8 | 6286 | 514.4 | 12903 | 92.72 | 709.6 | 46.95 | 266 |
| Mean (SD) | 41 (9) | | | 772 | 4527 | 412 | 5327 | 44 | 1014 | 46 | 303 |

| BNT162b2, COVID-19-Experienced | | | | | | | | | | | |
|---|---|---|---|---|---|---|---|---|---|---|---|
| | | | | D614G | | Delta | | Omicron BA.1 | | Omicron BA.2 | |
| Donor | Age | Sex | Comorbidities | 1 month post vax-2 | 1 month post booster | 1 month post vax-2 | 1 month post booster | 1 month post vax-2 | 1 month post booster | 1 month post vax-2 | 1 month post booster |
| 1 | 45 | M | Asthma | 5958 | 10443 | 5883 | 11917 | 169 | 1695 | 134.7 | 1456 |
| 2 | 37 | F | None | 8016 | 8991 | 5506 | 8828 | 208 | 1389 | 309.9 | 516.5 |
| 3 | 54 | F | Hypertension, Obesity | 11024 | 10453 | 2128 | 7968 | 174.3 | 1012 | 349.5 | 729 |
| 4 | 54 | M | Cardiovascular disease | 3989 | 7811 | 1406 | 8946 | 223.5 | 1566 | 79.65 | 1455 |

| | | | | | | | | | | | |
|---|---|---|---|---|---|---|---|---|---|---|---|
| 5 | 25 | F | None | 7768 | 10831 | 2073 | 8626 | 109.2 | 329.1 | 234.3 | 690.9 |
| 6 | 42 | F | Diabetes, Herpes simplex | 10747 | 9331 | 3240 | 17502 | 345.1 | 1364 | 180.5 | 712.2 |
| 7 | 43 | M | None | 5974 | 13891 | 2064 | 15888 | 171.7 | 334.3 | 211.7 | 346.1 |
| Mean (SD) | 41 (9) | | | 7639 | 10250 | 3185 | 11382 | 200 | 1098 | 214 | 843 |
